# CD47 predominates over CD24 as a macrophage immune checkpoint in cancer

**DOI:** 10.1101/2024.11.25.625185

**Authors:** Juliet Allen, Anna Meglan, Kyle Vaccaro, José Velarde, Victor Chen, Juliano Ribeiro, Jasmine Blandin, Sumeet Gupta, Ranjan Mishra, Raymond Ho, Jennifer Love, Ferenc Reinhardt, George W. Bell, Jin Chen, Robert Weinberg, Dian Yang, Jonathan Weissman, Kipp Weiskopf

## Abstract

Macrophages hold tremendous promise as effectors of cancer immunotherapy, but the best strategies to provoke these cells to attack tumors remain unknown. Here, we evaluated the therapeutic potential of targeting two distinct macrophage immune checkpoints: CD47 and CD24. We found that antibodies targeting these antigens could elicit maximal levels of phagocytosis when combined together in vitro. However, to our surprise, via unbiased genome-wide CRISPR screens, we found that CD24 primarily acts as a target of opsonization rather than an immune checkpoint. In a series of in vitro and in vivo genetic validation studies, we found that CD24 was neither necessary nor sufficient to protect cancer cells from macrophage phagocytosis in most mouse and human tumor models. Instead, anti-CD24 antibodies exhibit robust Fc-dependent activity, and as a consequence, they cause significant on-target hematologic toxicity in mice. To overcome these challenges and leverage our findings for therapeutic purposes, we engineered a collection of 77 novel bispecific antibodies that bind to a tumor antigen with one arm and engage macrophages with the second arm. We discovered multiple novel bispecifics that maximally activate macrophage-mediated cytotoxicity and reduce binding to healthy blood cells, including bispecifics targeting macrophage immune checkpoint molecules in combination with EGFR, TROP2, and CD71. Overall, our findings indicate that CD47 predominates over CD24 as a macrophage immune checkpoint in cancer, and that the novel bispecifics we created may be optimal immunotherapies to direct myeloid cells to eradicate solid tumors.

## INTRODUCTION

Macrophages are often the most common infiltrating immune cells in tumors, and they can be provoked to attack cancer under certain conditions (1–4). Just like T cells, macrophages are regulated by immune checkpoints that constrain their ability to recognize cancer cells as foreign and eliminate them (2–5). To date, the best-characterized macrophage checkpoint is the CD47/SIRPa interaction (2). CD47 is a surface antigen that is expressed on many cancers, and it functions by binding the inhibitory receptor, SIRPa, expressed on macrophages (2). Multiple drugs targeting this interaction have shown efficacy in preclinical models of both solid tumors and hematologic malignancies (6–9). Clinical trials using CD47-blocking therapies are ongoing and show promise for multiple types of cancer (10), indicating that activation of macrophage-mediated cytotoxicity can be an effective therapeutic strategy.

Based on the promise of targeting the CD47/SIRPa interaction, research efforts have focused on identifying additional macrophage immune checkpoints, or “don’t eat me” signals. Another surface antigen on cancer cells, CD24, is thought to act in this manner. In small cell lung cancer, the combination of CD47 blockade with anti-CD24 antibodies was found to enhance macrophage phagocytosis (9). A subsequent study proposed that CD24 on cancer cells functions by transmitting a direct inhibitory signal to macrophages, suggesting it acts as a macrophage immune checkpoint (11). However, the conservation of CD24 as a macrophage checkpoint across different cancers is unclear since unbiased genomic screens did not identify CD24 as a robust hit across multiple different human cancer types (12).

CD24 is a short, GPI-linked cell surface protein that is heavily glycosylated (13). It has been reported to interact with a number of ligands, including Siglec-10 (Siglec-G in mice), which can transduce inhibitory signals to some immune cell populations (14). In certain contexts, CD24 may also play an inhibitory role in B cell development (15). In cancer, prior studies examining the therapeutic potential of targeting CD24 have primarily relied upon xenograft models, which are limited by a lack of an adaptive immune system (11,16). Furthermore, the anti-CD24 antibodies used in these studies target human CD24 and do not cross-react with the murine CD24 homolog. Therefore, these models do not accurately reflect the therapeutic window for targeting CD24 since they underestimate toxicity and overrepresent selectivity to the tumor microenvironment. These pharmacologic properties can be better evaluated using syngeneic, immunocompetent mouse models. In the present study, we aimed to address these shortcomings by evaluating the therapeutic potential of targeting CD24 in immunocompetent mouse models of cancer. We found that dual-targeting of CD47 and CD24 could indeed elicit maximal levels of macrophage phagocytosis. However, via a series of genetic and pharmacologic experiments, we demonstrate that CD24 predominantly acts as a target of opsonization rather than a macrophage immune checkpoint. These findings provide insight into therapeutic targeting of CD24 in patients with cancer, and guided the development of novel bispecific antibodies that maximally activate macrophage anti-tumor functions.

## RESULTS

### Dual-targeting of CD47 and CD24 maximizes anti-tumor responses by macrophages

To compare expression of CD47 and CD24 in the murine system, we first performed cell surface profiling of seven different murine cancer cell lines (**Fig. 1A**). We found that the cell lines exhibited variable levels of expression of both CD47 and CD24, with no significant correlation between the expression of these two molecules on the cell surface (**Fig. 1B**). We found that 4 out of 7 lines expressed both CD47 and CD24, whereas 3LL ΔNRAS and MC38 cells expressed only CD47, and EL4 was negative for both CD47 and CD24.

**Figure 1:**
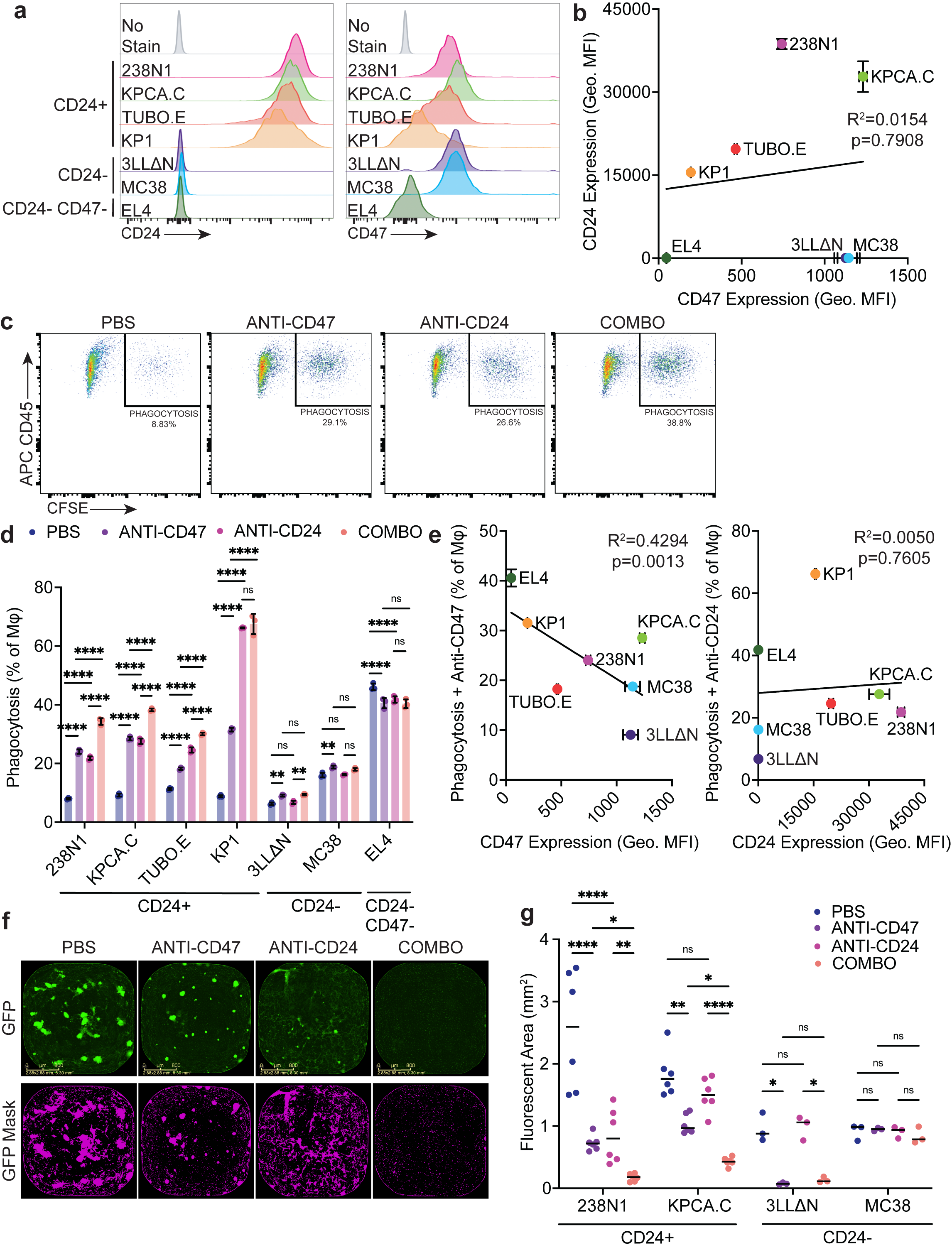
Anti-CD24 antibodies stimulate macrophage phagocytosis of mouse cancer cell lines. **A,** Histograms depicting cell surface expression of CD24 and CD47 by flow cytometry on mouse cancer cell lines. **B,** Correlation of CD24 and CD47 surface expression of cell lines shown in **A** by geometric MFI. Data shown as mean ± SD of 3 technical replicates. Simple linear regression was performed to assess correlation. **C,** Representative plots showing quantification of CD45+ phagocytic primary mouse macrophages co-cultured with CFSE+ KPCA.C. Co-cultures were exposed to vehicle control (PBS) or 10 ug/ml of monoclonal antibodies against mouse CD47, CD24, or the combination for 2 hours. Phagocytosis is represented as CD45+ macrophages that had engulfed CFSE+ KPCA.C cells as a percentage of the total macrophage population. **D,** Quantification of phagocytosis for cell lines in **A**. Cell lines are organized based on expression levels of each surface marker. Data represent mean ± SD of 3 technical replicates. **E,** Correlation of cell surface expression levels of CD47 and CD24 compared to phagocytosis upon treatment with the corresponding antibodies for each cell line. Data points depict mean ± SD from 3 replicates for each experiment. Correlation was assessed by simple linear regression. **F,** Representative microscopy images of GFP+ KPCA.C cells when co-cultured with primary mouse macrophages upon treatment with vehicle control (PBS), 10 ug/mL anti-CD47, 10 ug/mL anti-CD24, or the combination for 6.5 days. Top row depicts raw images of GFP+ fluorescence. Bottom row depicts purple GFP+ mask for above images used for quantification of cancer cell growth. Scale bar, 800 µm. **G,** Quantification of fluorescent well area from co-culture experiments for multiple cell lines after 6.5 days, organized by surface expression of CD24. Cancer cells were quantified by either green (KPCA.C, 3LL ΔNRAS, MC38) or red (238N1) fluorescent area based on their fluorophore expression. Data and means shown from one (3LL ΔNRAS, MC38) or two (238N1, KPCA.C) independent experiments with 3 technical replicates per experiment. **D,G,** statistical significance ns, *p<0.05, **p<0.01, ***p<0.001, ****p<0.0001 determined by two-way ANOVA with Holm-Sidak multiple comparison test.

To understand whether CD47 and CD24 levels could influence macrophage responses, we next performed in vitro phagocytosis assays using this panel of murine cancer cell lines. We co-cultured primary mouse macrophages with CFSE-labeled cancer cells for 2 hours with control, anti-CD47, or anti-CD24 therapy, then we measured phagocytosis by flow cytometry (**Fig. 1C**). We found that EL4, which lacks both CD47 and CD24 expression, had a relatively high baseline level of phagocytosis, but as expected, did not respond to antibodies targeting CD47 or CD24 (**Fig. 1D**). MC38 and 3LL ΔNRAS both exhibited minimal baseline levels of phagocytosis with small but significant increases upon anti-CD47 therapy (**Fig. 1D**). In contrast, all of the cell lines that are positive for CD24 surface expression (KPCA, 238N1, KP1, TUBO-EGFR) exhibited significant increases in phagocytosis with either anti-CD47 or anti-CD24 antibodies (**Fig. 1C and D**). The combination of these two antibodies further enhanced macrophage phagocytosis. In general, greater cell-surface CD47 expression inversely correlated with the response to anti-CD47 therapy (**Fig. 1E**), whereas CD24 expression did not correlate with phagocytosis upon anti-CD24 treatment (**Fig. 1E**).

We next investigated whether anti-CD47 or anti-CD24 therapy could be effective over longer periods of time. We employed co-culture assays in which macrophages and cancer cells are co-cultured for up to 7 days (17). These experiments integrate multiple mechanisms of macrophage-induced cell death, and they better model the long-term interactions between macrophages and cancer cells in the tumor microenvironment. Using this assay, we tested two murine cancer cell lines (KPCA.C and 238N1) that express both CD47 and CD24. We found that antibodies targeting either of these antigens could induce a moderate degree of macrophage-mediated cytotoxicity when used as single agents (**Fig. 1F and G**). Consistent with our phagocytosis assays, the combination of both anti-CD47 and anti-CD24 yielded remarkable anti-tumor responses, with elimination of nearly all cancer cells. As expected, cell lines that lacked CD24 surface expression (3LL ΔNRAS, MC38) exhibited no significant increase in macrophage cytotoxicity as a result of treatment with the anti-CD24 antibody (**Fig. 1G**). Together, these data suggest that dual-targeting of CD24 and CD47 can induce a robust anti-tumor response by macrophages.

### A genome-wide CRISPR screen indicates CD24 primarily functions as a target of opsonization

To understand which cancer cell genes regulate the ability of macrophages to respond to anti-CD24 therapies, we developed an intercellular CRISPR screening platform. We generated a variant of 238N1 lung adenocarcinoma cells that express Cas9 for gene knockout, and then introduced a pooled genome-wide sgRNA library (Gouda) into them. We expanded and selected the transduced cells, then subjected them to co-culture with primary mouse macrophages as a selective pressure. We performed co-culture for a period of 4 days under three distinct conditions: (i) cancer cells alone, (ii) cancer cells and macrophages, (iii) cancer cells, macrophages, and an anti-CD24 antibody (**Fig. 2A**). The surviving cancer cells were collected, and sgRNA representations and gene enrichment were evaluated. To our surprise, in the context of treatment with an anti-CD24 antibody, *Cd24a* was identified as one of the most significantly enriched hits, suggesting it is an “Eat me” signal expressed by the cancer cells in this context (**Fig. 2B, Supplementary Fig. S1**). Additionally, an extensive number of genes and processes involved in the biosynthesis of GPI-linked proteins were also enriched in our screen, functioning as “Eat me” signals upon treatment with anti-CD24 (**Fig. 2C and D, Supplementary Fig. S1**). Since CD24 is a glycophosphatidylinositol (GPI)-linked protein, these genes are required for post-translational processing that allows CD24 to be displayed on the cell membrane. Indeed, in validation studies, we found that knockout of one representative gene (Dpm1) could abrogate CD24 expression on the cell surface (**Fig. 2E**). The “Eat me” signals identified from our screen are associated with all aspects of GPI biosynthesis, including anchor biosynthesis within the endoplasmic reticulum and remodeling in the Golgi (**Fig. 2F**). These findings suggest the anti-CD24 antibody may function by opsonizing cancer cells and engaging Fc receptors on macrophages rather than disrupting an immune checkpoint.

**Figure 2:**
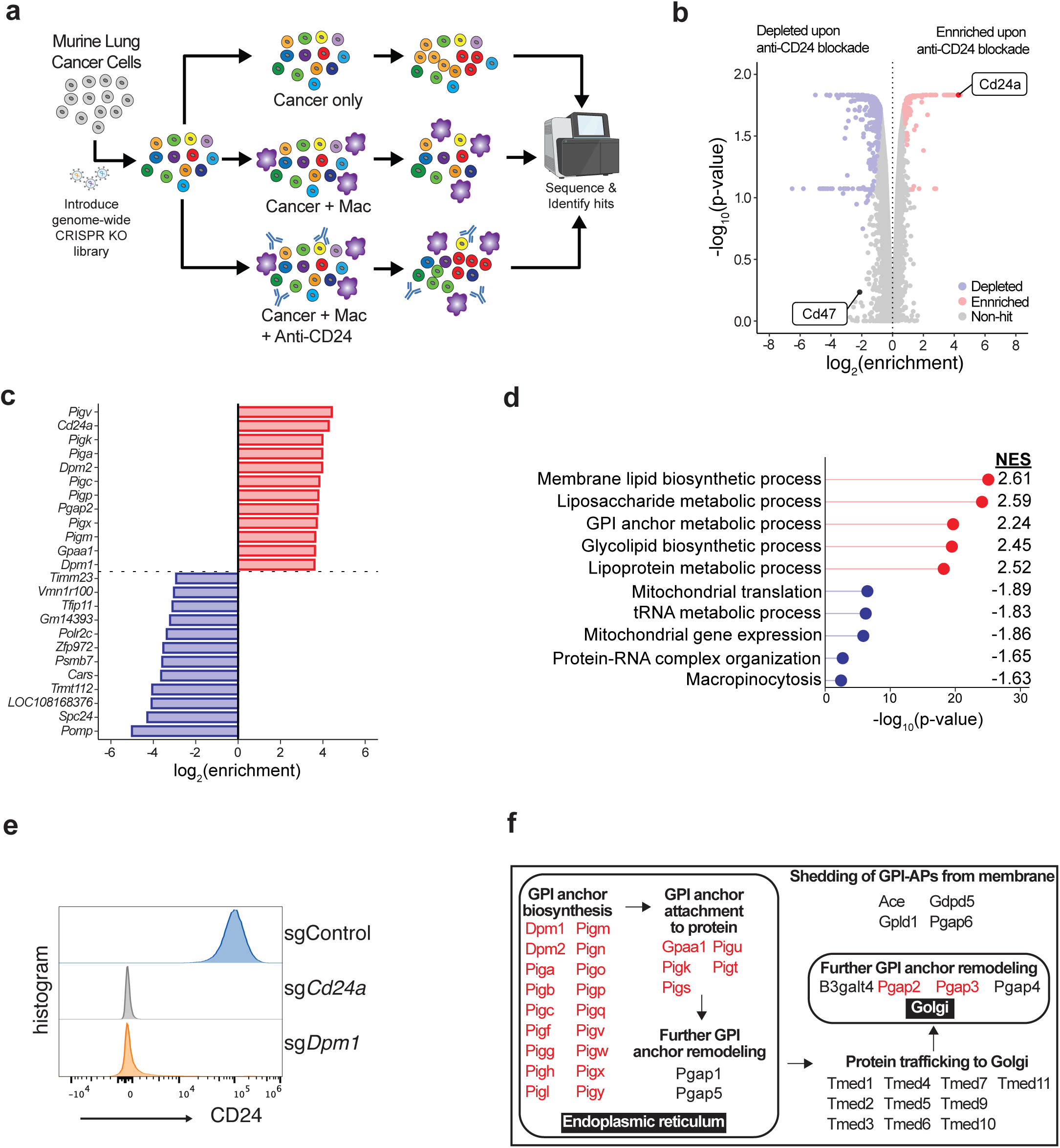
An unbiased, genome-wide CRISPR screen identifies CD24 as a target of opsonization rather than a macrophage immune checkpoint. **A,** Schematic of a genome-wide CRISPR screening strategy in murine lung cancer cell line 238N1 identifying differential responses to macrophage co-culture and treatment with anti-CD24. **B,** Volcano plot of genome-wide screen comparing treatment with macrophages and anti-CD24 against treatment with macrophages alone. Genes with a discovery score > 5 (FDR < 0.05) are highlighted, with enriched genes shown in red and depleted genes shown in blue. **C,** Top differentially enriched and depleted genes with respective log_2_(enrichment) from differential enrichment analysis depicted in **B. D,** Differentially enriched GO Biological Process pathways comparing treatment with macrophages and anti-CD24 against treatment with macrophages alone. Top 5 enriched and depleted pathways are shown (p-adjusted <0.05). **E,** Analysis of CD24 expression on 238N1 cells by flow cytometry upon treatment with sgRNAs targeting the indicated genes. **F,** Schematic of genes involved in CD24 biosynthesis and regulation of cell-surface display. Gene hits from the screen depicted in **B** and **C** are highlighted in red.

### Genetic analysis demonstrates CD24 does not protect cancer cells from macrophage attack in vitro

To further understand the functional effects of targeting CD47 or CD24, we examined genetic perturbations of these genes in KPCA.C and 238N1 cells. We ablated CD47 or CD24 expression in each line by either CRISPRi (238N1) or CRISPR knockout (KPCA.C) (**Fig. 3A**). For KPCA.C cells, we also generated a double-knockout (DKO) variant in which both genes were genetically ablated. We tested these cell lines in phagocytosis assays using primary mouse macrophages. For both cell lines, we observed a significant increase in phagocytosis as a consequence of CD47 ablation, and this effect was dramatically enhanced by the addition of an anti-CD24 antibody (**Fig. 3B-D**). These results are consistent with the known function of CD47 and its role in sensitizing to tumor-opsonizing antibodies that engage Fc receptors on macrophages (18). In contrast, CD24 ablation resulted in no significant increase in phagocytosis at baseline, and it abrogated the effects of the anti-CD24 antibody (**Fig. 3B-D**). Furthermore, when using the KPCA.C DKO cell line, no further enhancement was observed in phagocytosis above what was seen with CD47 knockout alone. These data again suggest that CD24 is not required to protect cancer cells from macrophage phagocytosis in vitro.

**Figure 3:**
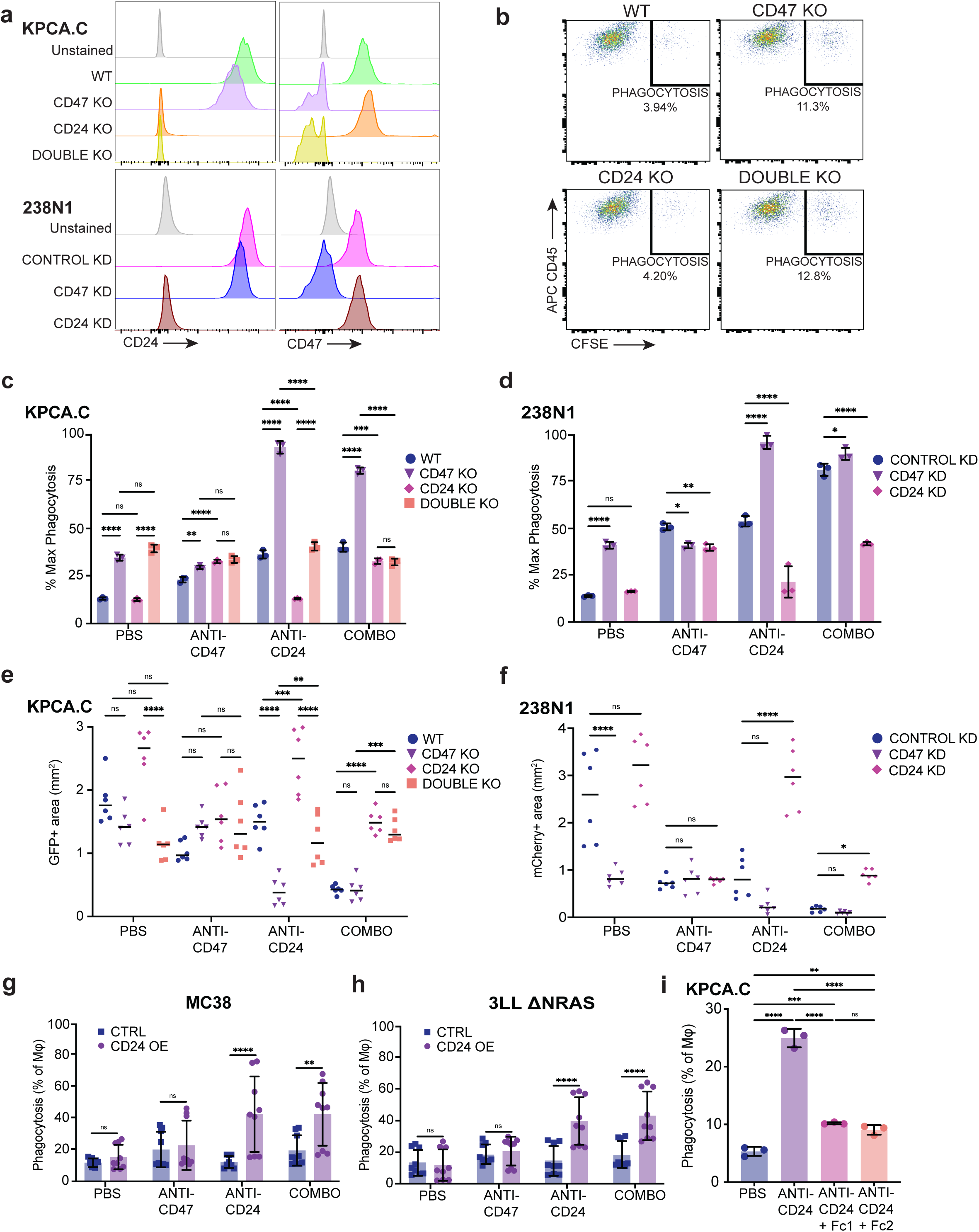
Genetic ablation of CD24 does not influence macrophage anti-tumor responses in vitro. **A,** Representative histograms demonstrating cell surface expression of CD47 and CD24 on knockouts of KPCA.C and knockdowns of 238N1 by flow cytometry. **B,** Representative gating of phagocytic APC CD45+ mouse macrophages when co-cultured with the indicated CFSE+ KPCA.C knockouts treated with vehicle control (PBS) for 2 hours. Phagocytic macrophages are calculated as CD45+ cells that have engulfed CFSE+ cancer cells after 2 hours as a percent of all macrophages. **C,D,** Quantification of phagocytosis as a percentage of the maximum phagocytic response of macrophages using KPCA.C knockout cells (**C**) or 238N1 knockdown cells (**D**) treated with vehicle control (PBS), anti-mouse CD47 antibody, anti-mouse CD24 antibody, or the combination. Data represents mean ± SD of 3 technical replicates. **E,F,** Quantification of fluorescent well area as a measure of GFP+ KPCA.C knockout cells (**E**) or mCherry+ 238N1 knockdown cells (**F**) growth after co-culture with primary mouse macrophages and the indicated antibodies on day 6.5. Data represents mean ± SD from two independent experiments of 3 technical replicates each. **G,H,** Quantification of phagocytosis using CFSE+ MC38 (**G**) or 3LL ΔNRAS (**H**) cancer cells that overexpress CD24 after co-culture with primary mouse macrophages and the indicated antibodies. Data represent mean ± SD from 3 individual experiments each containing 3 technical replicates. **I,** Quantification of phagocytosis using StayGold+ KPCA.C cancer cells treated with vehicle control (PBS), or anti-mouse CD24 antibody, in the absence or presence of FcR blocking reagents (Fc1, anti-mouse Truestain clone 93; Fc2, anti-mouse CD16/CD32 clone 2.4G2). Data represents mean ± SD of 3 technical replicates. (**C-H**) ns, *p<0.05, **p<0.01, ***p<0.001, ****p<0.0001 by two-way ANOVA with Holm-Sidak multiple comparison test.

Similarly, we also performed long-term co-culture assays with the knockout and knockdown lines. CD47 knockdown significantly decreased 238N1 cell growth in co-culture with primary mouse macrophages, and it sensitized both KPCA.C and 238N1 cells to further macrophage-mediated cytotoxicity in the presence of an anti-CD24 antibody, mirroring the results of our phagocytosis assays (**Fig. 3E and F**). However, CD24 ablation demonstrated no significant inhibition of tumor growth for either cell line at baseline or in combination with anti-CD47 (**Fig. 3E and F)**. Moreover, upon treatment with an anti-CD24 antibody, the cell lines lacking CD24 expression were resistant to the effects of the antibody and grew unhindered and comparable to untreated cells (**Fig. 3E and F**). Again the DKO cell line exhibited no enhanced sensitivity relative to CD47 knockout alone. Together, these results further implicate CD24 as a target of opsonization.

To understand whether CD24 expression is sufficient to protect against macrophage phagocytosis, we next investigated the effects of overexpressing CD24 using MC38 and 3LL ΔNRAS cells, which lack native CD24 expression (**Supplementary Fig. S2**). In both models, compared to control cells, CD24 overexpression did not significantly inhibit phagocytosis in control or antibody treatment conditions (**Fig. 3G and H**). Instead, treatment with an anti-CD24 antibody significantly enhanced phagocytosis for both CD24 overexpression lines. These data indicate that CD24 expression is not sufficient to protect cancer cells from macrophage attack. To directly evaluate dependency on Fc receptors (FcRs), we examined the ability of the anti-CD24 antibody to induce phagocytosis in the absence or presence of FcR blocking reagents (**Fig. 3I**). We found that in the presence of FcR blockade, the phagocytosis stimulating abilities of the antibody were significantly abrogated. Together, these findings further indicate that CD24 primarily functions as a target of opsonization.

### CD47 predominates over CD24 as a macrophage checkpoint in syngeneic tumor models

Since our in vitro experiments may not fully recapitulate the state of macrophages within the tumor microenvironment, we evaluated the CD24 perturbation lines in vivo. First, we engrafted wild-type, CD47 knockout, CD24 knockout, or DKO variants of KPCA.C cells subcutaneously into the flanks of C57BL/6 mice and measured tumor volumes over time (**Fig. 4A**). We observed modest inhibition of tumor growth with CD24 knockout, but both the CD47 knockout and DKO lines grew significantly slower than the CD24 knockout line. There was no significant difference between the CD47 knockout and DKO lines (**Fig. 4A**). The growth trends of these cell lines were similar when the KPCA.C variants were engrafted into NSG mice, which lack an adaptive immune system (**Fig. 4B**), suggesting any anti-tumor immune response is due to myeloid cells. To understand if the site of engraftment may influence the response to these cell lines, we also engrafted KPCA.C cells directly into the peritoneal cavities of mice. This site models advanced stages of disease and is enriched with resident macrophages (19–21). We found that CD47 knockout significantly prolonged survival of mice engrafted with intraperitoneal tumors, whereas no significant difference was observed between control KPCA.C cells and CD24 knockout cells (**Fig. 4C**). Together, these findings indicate that CD47 predominates over CD24 as a macrophage immune checkpoint in the KPCA.C model.

**Figure 4:**
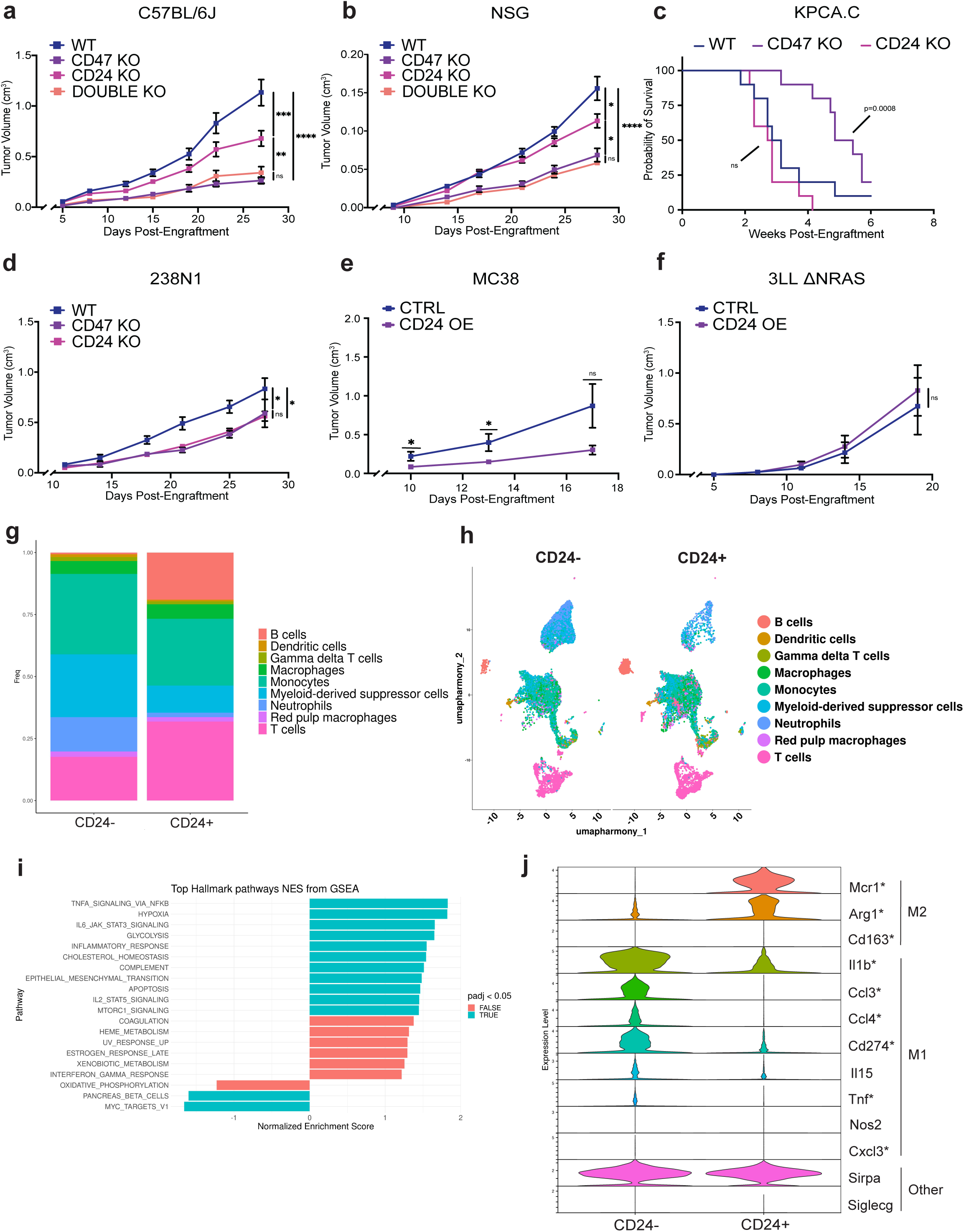
CD24 exhibits limited activity as a macrophage immune checkpoint in immunocompetent tumor models. **A,** Tumor volume growth curve for KPCA.C cells with the indicated knockout engrafted subcutaneously into C57BL/6J mice. Data represents mean ± SEM for a total of n = 20 mice per cohort with data combined from two independent experiments. **B,** Tumor volumes for KPCA.C knockout cells engrafted subcutaneously in NSG mice. Data represents mean ± SEM for n = 10 mice per cohort. **C,** Kaplan-Meier curve for C57BL/6J mice engrafted intraperitoneally with KPCA.C cells with the indicated knockouts. Data represent survival analysis for n = 10 mice per group. **D,** Tumor volume growth curve for 238N1 cells with the indicated knockouts engrafted subcutaneously into B6129SF1/J mice. Data represent mean ± SEM from 3 independent experiments each with n = 5 mice per cohort. **E,F,** Tumor volumes for MC38 (**E**) or 3LL ΔNRAS (**F**) variants overexpressing CD24 that were engrafted subcutaneously into C57BL/6J mice. Data represent mean ± SEM from one (**F**) or two (**E**) independent experiments each containing n = 5 mice per group. **G,** Relative frequencies of CD45+ immune cells sorted from CD24-negative versus CD24-positive tumors as evaluated by single-cell RNA-sequencing analysis. **H,** UMAP showing identified clusters from immune cells sorted from CD24-negative versus CD24-positive tumors. **I,** Gene enrichment analysis of immune cells from CD24-negative versus CD24-positive tumors. **J,** Violin plot showing differentially expressed M1-like versus M2-like genes within the myeloid-derived suppressor cell population. *adjusted p value <0.05. (**A-F**) ns, *p<0.05, **p<0.01, ***p<0.001, ****p<0.0001 by one-way ANOVA with Holm-Sidak multiple comparison test (**A,B,D-F**) or by Mantel-Cox log-rank test (**C**) at the indicated endpoint.

We performed similar in vivo experiments using 238N1 cells. We engrafted wild-type, CD47 knockdown, and CD24 knockdown variants of 238N1 into the flanks of immunocompetent mice, and we measured tumor volumes over time. We observed that CD47 knockdown or CD24 knockdown only caused modest inhibition of tumor growth in this model (**Fig. 4D**), suggesting that there can be differential sensitivity to targeting macrophage immune checkpoints in vivo.

We also examined whether overexpression of CD24 could be sufficient to protect cancer cells from macrophage attack in vivo. Using both MC38 and 3LL ΔNRAS cells, we compared engraftment of control cells versus CD24 overexpression cells. We found that CD24 overexpression did not significantly enhance the growth of tumors (**Fig. 4E and F**), indicating that CD24 is not sufficient to protect cancer cells from macrophage-mediated cytotoxicity in vivo.

Although CD24 perturbations may have only modest effects on regulating tumor growth in vivo, we considered whether these perturbations could alter immune activation within the tumor microenvironment. We collected tumors from KPCA.C, 238N1, and MC38 experiments, dissociated them to single cell suspensions, sorted CD45+ immune cells, and subjected them to single-cell RNA-sequencing (scRNA-seq). We compared CD24-tumors (KPCA.C CD24 knockout, 238N1 CD24 knockout, and MC38 control) to CD24+ tumors (KPCA.C control, 238N1 control, and MC38 CD24 overexpression). We observed a greater degree of myeloid cell infiltration in CD24-tumors, including increases in neutrophils and myeloid-derived suppressor cells (**Fig. 4G and H**). In contrast, we observed a greater degree of lymphoid infiltration in CD24+ tumors, including B cells and T cells (**Fig. 4G and H**). By gene set enrichment analysis (GSEA), we found that immune cells from CD24-tumors exhibited enrichment for expression programs associated with TNFa signaling, hypoxia, IL6 signaling, glycolysis, and an inflammatory response (**Fig. 4I**). These transcriptional programs were similarly activated by loss of CD47 in the cancer cells (**Supplementary Fig. S3**). We observed some of the greatest differences within the myeloid-derived suppressor cell (MDSC) population in CD24-tumors, where “M1-like” genes were enriched and “M2-like” genes were depleted (**Fig. 4J**), consistent with MDSC repolarization towards a pro-inflammatory state (22). Together, these findings suggest that genetic perturbations of CD24 alone may not substantially influence tumor growth, but may nevertheless alter immune infiltrates and activation states within the tumor microenvironment.

### Therapeutic targeting of CD24 in mice is limited by hematologic toxicity

Regardless of the underlying mechanism, we unambiguously observed that anti-CD24 antibodies are capable of stimulating robust macrophage-mediated cytotoxicity in vitro. Therefore, we sought to determine the therapeutic potential of anti-CD24 antibodies in a fully immunocompetent mouse model. We engrafted mice with wild-type, CD47 knockout, or CD24 knockout KPCA.C cells. For each variant, we randomized the mice into two treatment groups in which mice were treated with vehicle control or an anti-CD24 antibody. Again, we observed that CD47 knockout significantly inhibited the growth of KPCA.C tumors, whereas minimal growth inhibition was observed with CD24 knockout cells (**Fig. 5A**). We found that the anti-CD24 antibody had no significant effect on tumor growth for any of the KPCA.C variants (**Fig. 5A**). Unexpectedly, two mice treated with the anti-CD24 antibody exhibited acute toxicity as a result of the treatment, suggesting that anti-CD24 therapy may be limited by a low therapeutic index.

**Figure 5:**
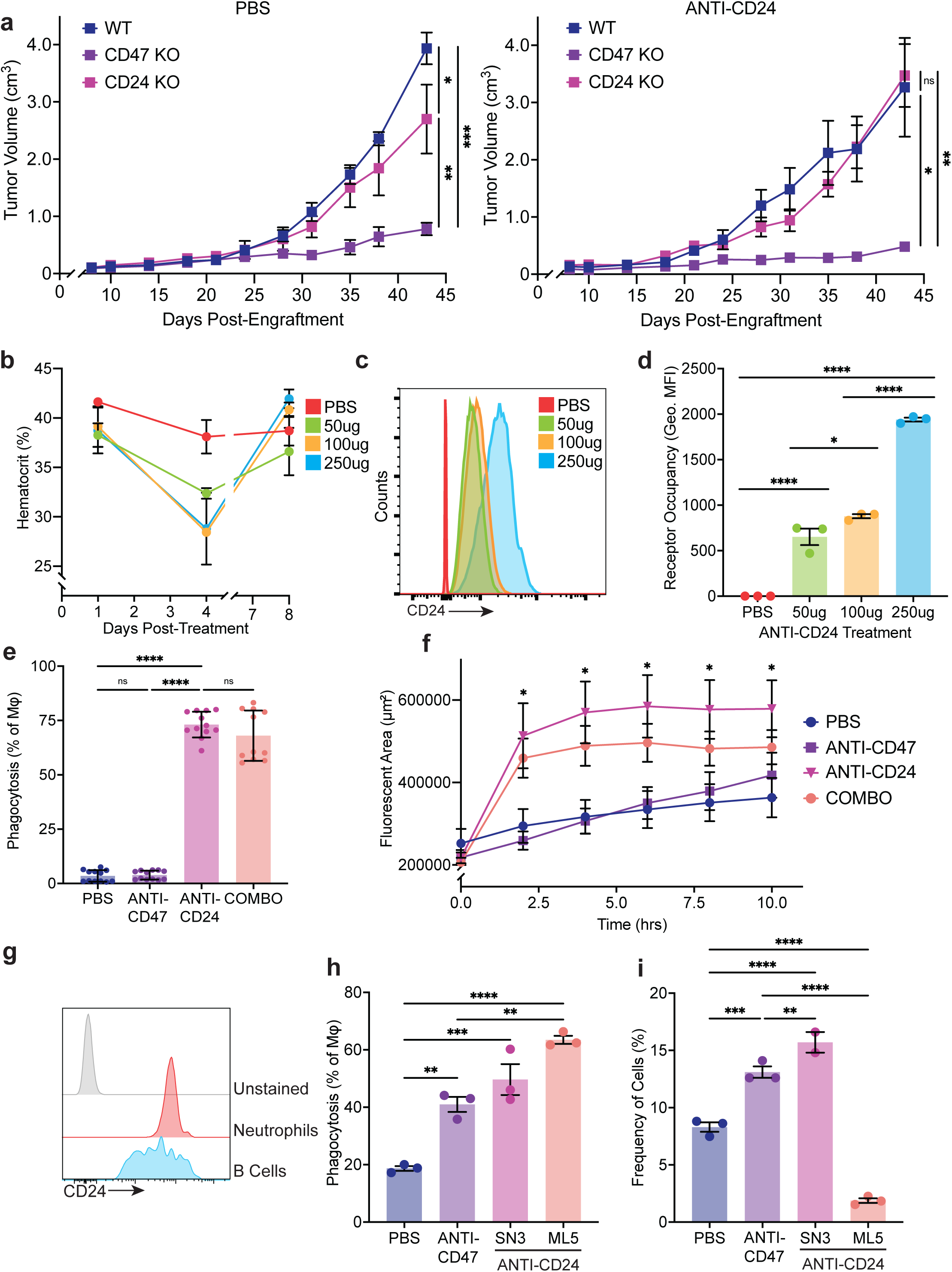
Therapeutic targeting of CD24 poses risks of hematologic toxicity. **A,** Tumor volume curves of KPCA.C variants engrafted into C57BL/6J mice treated with vehicle control (PBS) or an anti-mouse CD24 antibody. Data represents mean ± SEM of n = 5 mice per group. **B,** C57BL/6J mice were treated with PBS control or 50ug, 100ug, or 250ug of an anti-mouse CD24 antibody. Hematocrit levels are represented as the mean ± SEM of 3 mice per group measured at the indicated days post-treatment. **C,** Representative histogram depicting receptor occupancy of anti-mouse CD24 antibody on the surface of mouse red blood cells as detected by staining for antibody binding. **D,** Quantification of receptor occupancy of anti-CD24 on the surface of red blood cells. Data represents mean ± SEM of n = 3 mice per group. **E,** Mouse red blood cells were isolated from C57BL/6J mice, labeled with CFSE+ and co-cultured with primary mouse macrophages for 2 hours under treatment with PBS control or anti-mouse CD24 antibody. Phagocytosis is represented as the percentage of total macrophages engulfing CFSE+ red blood cells. Data represent mean ± SD of 12 technical replicates from two independent experiments. **F,** Phagocytosis of pHrodo+ mouse red blood cells over time by primary mouse macrophages treated with the indicated therapies. pHrodo+ area was measured over time, and data represent mean ± SEM of 3 technical replicates. **G,** CD24+ surface expression profiling of human whole blood showing CD15+ neutrophil and CD20+ B cell immune cell populations by flow cytometry. Histograms show representative example from 3 technical replicates. **H,** Phagocytosis of CFSE+ human neutrophils by human macrophages following 2 hour co-culture with the indicated therapies. Phagocytosis is represented as a percentage of total macrophages. Data shown as mean ± SEM of 3 technical replicates. **I,** CD15+ neutrophil profiling as percentage of live cells from RBC depleted whole blood after 16 hours of culture in the presence of the indicated antibodies. Data depict mean ± SEM of 3 technical replicates. (**A,D,E,F,H,I**) ns, *p<0.05, **p<0.01, ***p<0.001, ****p<0.0001 by one-way ANOVA with Holm-Sidak multiple comparison test at the indicated endpoint.

Given the rapid onset of mortality as a consequence of anti-CD24 treatment, we hypothesized that the antibody may have acute effects on the hematologic system. Indeed, we found that treatment with a single dose of the anti-CD24 antibody caused an acute drop in the hematocrit of the mice, with a nadir at 3 days post-treatment (**Fig. 5B**). The anti-CD24 antibody bound to the surface of all red blood cells in treated mice (**Fig. 5C and D**), and the drop in hematocrit corresponded to receptor occupancy on the surface of red blood cells over time (**Fig. 5C and D, Supplementary Fig. S4**). The observed hematologic toxicity exhibited a dose-response relationship (**Fig. 5B**). To evaluate the mechanism of toxicity, we performed in vitro phagocytosis assays with mouse macrophages and syngeneic red blood cells. We found the anti-CD24 antibody induced substantial phagocytosis of red blood cells in vitro (**Fig. 5E and F**). These findings suggest that anti-CD24 antibodies may carry a significant risk for hematologic toxicity.

### Therapeutic targeting of CD24 poses a risk for neutrophil toxicity in the human system

To understand whether similar side effects may occur with CD24-targeting therapies in humans, we analyzed blood specimens from healthy human donors. We found that CD24 exhibited a different expression pattern in mice versus humans: CD24 was not present on the surface of human red blood cells, but instead was highly expressed on neutrophils and B cells (**Fig. 5G**). To understand whether anti-CD24 antibodies could cause toxicity to human immune cell subsets, we performed human macrophage phagocytosis assays using primary human neutrophils as targets. We tested two different anti-CD24 antibodies and compared them to an anti-CD47 antibody. We found that both anti-CD24 antibodies were able to stimulate significant phagocytosis that was comparable to or exceeded that of anti-CD47 (**Fig. 5H**). Additionally, we performed culture of unfractionated human leukocytes with anti-CD24 antibodies. We found that one of the anti-CD24 antibodies (clone ML5) caused substantial depletion of neutrophils in this system (**Fig. 5I**), likely due to its murine IgG2a isotype that can elicit greater immune effector functions. We also identified a proinflammatory cytokine profile in the supernatants that corresponded to neutrophil depletion in these samples, including secretion of IL-1b, MCP-3, and ENA-78 (**Supplementary Fig. S5**). These findings suggest anti-CD24 therapies pose risks of on-target toxicity to human neutrophils that should be cautiously monitored in clinical trials testing anti-CD24 antibodies.

### Genetic perturbations demonstrate that CD24 is not a universal macrophage immune checkpoint in human cancers

Since the effects of CD24 perturbations have only been studied in a limited number of models, we examined CD24 ablation in additional human cancer cell lines. We identified four cell lines that express varying degrees of cell-surface CD24 (NCI-H3122, NCI-H358, DLD-1, NIH-OVCAR3), and we generated knockdown or knockout variants of these cell lines using CRISPR-based approaches (**Supplementary Fig. S6**). We tested each of the lines in phagocytosis assays using primary human macrophages. As in our murine studies, we found that knockdown or knockout of CD24 did not enhance macrophage phagocytosis at baseline, in response to an anti-CD47 antibody, or in response to an anti-EGFR antibody (**Fig. 6A-D**). Conversely, knockout of CD47 sensitized cancer cells to phagocytosis in response to an anti-CD24 antibody (**Fig. 6C and D**), consistent with its ability to stimulate antibody-dependent phagocytosis.

**Figure 6:**
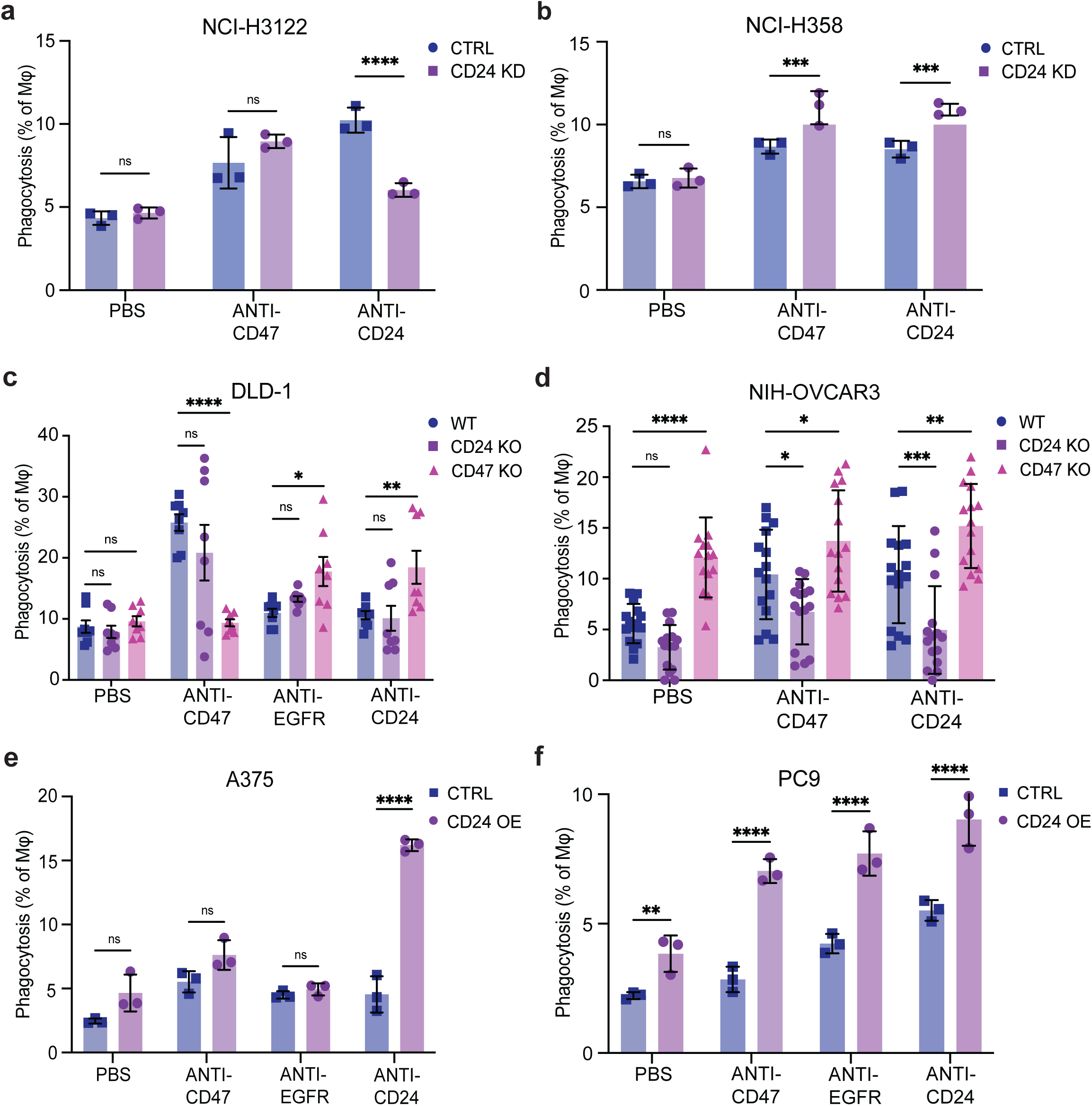
CD24 does not influence human macrophage phagocytosis in vitro. **A,B,** Phagocytosis assays using primary human macrophages and CD24 knockdown variants of NCI-H3122 (**A**), or NCI-H358 (**B)**. Data represent mean ± SD of 3 technical replicates from an individual human macrophage donor per cell line. **C,D,** Phagocytosis assays using human macrophages and CD24 knockout variants deriving from DLD-1 (**C**) or NIH-OVCAR3 (**D**). Data represent mean ± SD of 2 independent experiments containing 2 or 3 technical replicates each with a total of n = 3 individual human macrophage donors (**C**) or 3 independent experiments containing 3 technical replicates with a total of n = 5 individual human macrophage donors (**D**). **E,F,** Phagocytosis assays using CD24 overexpression lines deriving from A375 (**E**) or PC9 (**F**). Data represent mean ± SD of 3 technical replicates from an individual human macrophage donor per cell line. All phagocytosis was quantified as a percentage of total macrophages. (**A-F**) ns, *p<0.05, **p<0.01, ***p<0.001, ****p<0.0001 by one-way ANOVA with Holm-Sidak multiple comparison test.

To evaluate whether CD24 is sufficient to protect human cancer cells from phagocytosis, we also tested two cell lines that exhibit low or absent CD24 expression (A375 and PC9) and we overexpressed CD24 via lentiviral transduction (**Supplementary Fig. S6**). We found that CD24 overexpression did not decrease phagocytosis at baseline or in response to anti-CD47 (**Fig. 6E and F**). However, CD24 overexpression significantly sensitized the cancer cells to phagocytosis with an anti-CD24 antibody (**Fig. 6E and F**). Overall, these findings are consistent with the results from our murine studies and indicate CD24 primarily functions as a target of opsonization in the human system.

Since the macrophages in our in vitro studies may not fully model macrophages in the tumor microenvironment, we also engrafted the CD24 perturbation lines into NSG mice as xenograft models. NSG mice lack functional T, B, and NK cells, but contain macrophages and myeloid immune cells that can respond to macrophage-directed therapies in vivo (7,18,23). These models have been considered gold standards for evaluating macrophage-directed therapies in vivo. We engrafted three different CD24 knockdown lines in vivo: NCI-H358 cells, DLD-1 cells, NCI-H3122 cells (**Fig. 7A-C**). We only observed a significant but modest reduction in tumor growth from CD24 knockdown in the NCI-H3122 model (**Fig. 7A**). Genetic knockout of CD24 in NIH-OVCAR3 cells also had no significant effect on tumor growth (**Fig. 7D**). To understand whether CD24 could be sufficient to protect human cancer cells from macrophage attack in vivo, we also tested the two CD24 overexpression lines. In each case, no significant effect was observed from overexpression of CD24 on the cell surface (**Fig. 7E and F**). These studies suggest that for the majority of cancers, CD24 is neither necessary nor sufficient to protect cancer cells from macrophage phagocytosis.

**Figure 7:**
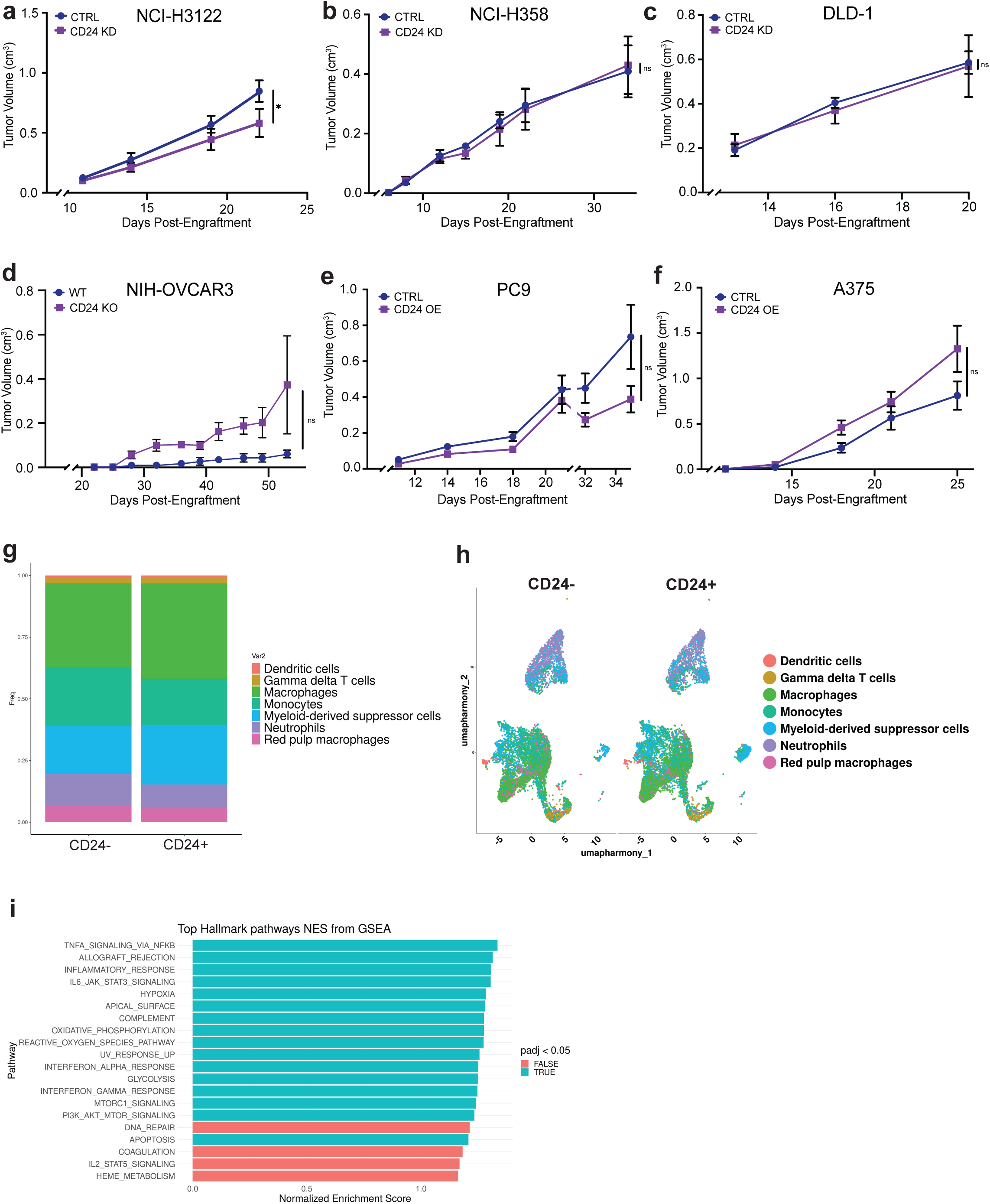
Perturbations of human CD24 minimally influence tumor growth in xenograft models but alter immune activation. **A-C,** Tumor volume growth curves of NSG mice engrafted subcutaneously with control or CD24 knockdown tumors deriving from NCI-H3122 (**A**), NCI-H358 (**B**), or DLD-1 (**C**). Data represents mean ± SEM from a total of n = 15 mice per cohort combined from 3 individual experiments (**A**), or one individual experiment (**B,C**) containing n = 5 mice per group. **D,** Tumor volume growth curves of NSG mice engrafted with control or CD24 knockout tumors deriving from NIH-OVCAR3 cells. Data represents mean ± SEM of n = 5 mice per group. **E,F,** Tumor volume growth curves of NSG mice engrafted with control or CD24 overexpression lines deriving from A375 (**E**) or PC9 (**F**). Data represents mean ± SEM of n = 5 mice per group. **G,** Relative frequencies of CD45+ immune cells sorted from CD24-negative versus CD24-positive tumors as evaluated by single-cell RNA-sequencing analysis. **H,** UMAP showing identified cell clusters from immune cells sorted from CD24-negative versus CD24-positive tumors. **I,** Gene set enrichment analysis of immune cells from CD24-negative versus CD24-positive tumors. (**A-F**) ns, *p<0.05 by two-tailed t test at the indicated endpoint.

As with our syngeneic tumor models, we also performed scRNA-seq from the mouse xenograft experiments to understand whether genetic perturbations in CD24 can alter immune activation. We compared sorted immune cells from CD24-tumors (NCI-H3122 CD24 knockdown, DLD-1 CD24 knockdown, and A375 control) and compared these to CD24+ tumors (NCI-H3122 control, DLD-1 control, or A375 CD24 overexpression). We observed that immune cell subsets were similar in quantity across the two groups (**Fig. 7G and H**). As with our syngeneic mouse models, GSEA revealed enrichment for expression of hallmark gene sets associated with TNFa and IL-6 signaling, an inflammatory response, and hypoxia (**Fig. 7I**).

These changes corresponded to polarization of myeloid cells towards an “M1-like” state in the CD24-tumors. Consistently, these findings indicate that although CD24 may influence immune cell activation within the tumor microenvironment, genetic perturbations of CD24 do not significantly affect the growth of tumors in the majority of models tested.

### Therapeutic Targeting of CD47 Predominates over CD24 in a Bispecific Format

To leverage our findings and further evaluate the therapeutic potential of targeting CD47 and CD24, we developed a compendium of novel bispecific antibodies. We reasoned that a bispecific format that targets CD47 or CD24 with one arm and a tumor-specific antigen with the other arm could enhance specificity to the tumor microenvironment and minimize toxicity to normal cells (**Fig. 8A**). We therefore created a library of scFvs and binding domains to a variety of tumor antigens (e.g., EGFR, HER2, TROP2, among others) and macrophage immune checkpoint molecules (CD47, CD24, SIRPa, and PD-1). We tested two different binding domains for targeting CD47: CV1 (a high-affinity binder) and WTa2d1 (a low-affinity binder) (18). We tested three different CD24-binding domains (CD24-1, CD24-2, CD24-3). We engineered each construct to be expressed as a human IgG1 knob-into-hole heterodimer (24). We crossed each macrophage checkpoint arm with each tumor antigen arm to create a total of 77 novel bispecific antibodies. We individually tested the functional anti-tumor effects of each of these antibodies in co-culture assays using primary human macrophages and fluorescent DLD-1 cells (**Fig. 8B-G**). We found that that majority of bispecific antibodies containing CV1, WTa2d1, or CD24-3 exhibited significant functional anti-tumor effects (**Fig. 8B-G**). CV1 exhibited the greatest anti-tumor effects, nearly eliminating all cancer cells across all formats (**Fig. 8C and F**). WTa2d1 exhibited robust anti-tumor efficacy when paired with an EGFR, TROP2, or CD71 binding arms (**Fig. 8D**). The CD24-3 bispecifics exhibited substantial anti-tumor effects when paired with most tumor-targeting arms (**Fig. 8E**).

**Figure 8:**
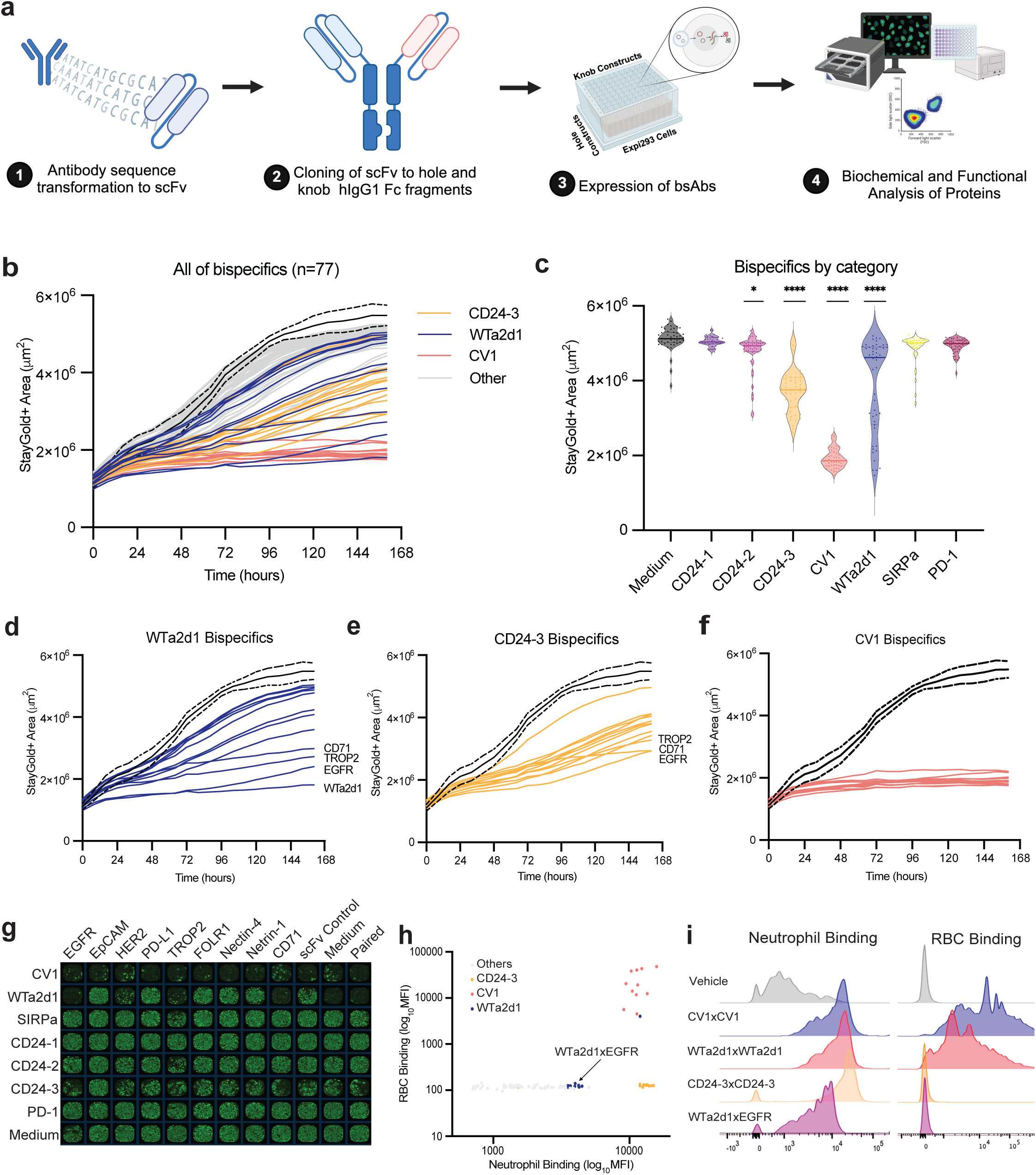
Bispecific antibodies can maximize anti-tumor responses by macrophages while minimizing binding to healthy cells. **A,** Diagram showing process for high-throughput development and functional evaluation of bispecific antibodies targeting macrophage immune checkpoints. Antibody sequences were transformed into scFvs and cloned into a knob-into-hole format using a human IgG1 Fc. Constructs targeting macrophage immune checkpoints (CD47, CD24, SIRPa, PD-1) were cloned into knob formats and crossed with tumor-binding constructs in a hole format. Bispecific antibodies (n = 77) were expressed in Expi293F cells and used for downstream biochemical and functional analysis. **B,** Growth of StayGold+ DLD-1 cells in co-culture with human macrophages and each bispecific antibody. Each curve represents the mean for an individual bispecific antibody from 4 replicates. Black curve with hashed lines represents mean and 95% CI of control wells**. C,** Anti-tumor efficacy of bispecific antibodies at approximately t = 6.5 days as evaluated by macrophage checkpoint category. *p<0.05, ****p<0.0001 by one-way ANOVA with Dunnett’s multiple comparisons test. **D-F,** Growth curves for each of the WTa2d1 constructs (**D**), CD24-3 constructs (**E**), or CV1 constructs (**F**). **G,** Representative whole-well imaging of co-cultures treated with different bispecific antibodies at approximately t = 6.5 day. Green signal depicts growth of StayGold+ DLD-1 cells. Rows contain different macrophage checkpoint arms, while columns contain different tumor-binding arms. **H,** Scatter plot showing binding of each bispecific antibody to human neutrophils versus red blood cells. **I,** Representative histograms showing binding of the indicated bispecific antibodies to human neutrophils and red blood cells.

An ideal bispecific antibody should not only exert a robust anti-tumor response, but should also avoid on-target toxicity to the hematologic system. We therefore determined whether any of the bispecific antibodies could decrease this risk. We found that all constructs containing CV1, the high-affinity CD47-binding domain, exhibited intense binding to red blood cells and neutrophils (**Fig. 8H and I**). CD24-3 did not bind to red blood cells, but intensely labeled neutrophils (**Fig. 8H and I**), again indicating the risk of hematologic toxicity for targeting CD24 in the human system. The WTa2d1 x WTa2d1 homodimer also exhibited substantial binding to red blood cells (**Fig. 8H and I**). In contrast, the heterodimeric WTa2d1 bispecifics exhibited significantly reduced binding to both red blood cells and neutrophils (**Fig. 8H and I**). Thus, bispecific antibodies including a WTa2d1 domain appear to be optimal to maximize the therapeutic index, particularly WTa2d1 x EGFR, WTa2d1 x TROP2, and WTa2d1 x CD71. In sum, these efforts indicate that CD47 again predominates over CD24 for therapeutic targeting and further illuminate the most effective strategies to maximize anti-tumor responses by macrophages.

## DISCUSSION

We have conducted a rigorous evaluation of CD47 and CD24 as macrophage immune checkpoints in cancer. Across mouse and human systems, we found that in the vast majority of cases, CD24 is neither necessary nor sufficient to protect cancer cells from macrophage attack. Instead, our findings indicate that anti-CD24 antibodies primarily act in an Fc-dependent manner to opsonize cancer cells, thereby stimulating macrophage phagocytosis. These conclusions are supported by multiple syngeneic mouse tumor models in vitro and in vivo, fully human systems in vitro, and xenograft tumor models. We consistently found that the effects of CD47 ablation were greater than genetic perturbations of CD24. Thus, our overall findings demonstrate that CD47 predominates over CD24 as a macrophage immune checkpoint in cancer.

The results of our unbiased screening methods are consistent with those of Kamber et al, who also did not identify CD24 as a robust immune checkpoint from genome-wide screens using a co-culture system of macrophages and human cancer cell lines (12). Our findings differ in part from those reported by Barkal et al., who described a putative role for CD24 in regulating anti-tumor responses by macrophages (11). Our study suggests that although CD24 may exhibit a modest function as a macrophage immune checkpoint, the predominant activity of antibodies targeting this antigen are driven by opsonization. We do not exclude the possibility that CD24 may function as a more substantive immune checkpoint for certain cancers, and this function could depend on factors such as the degree of CD24 expression, the presence of other “don’t eat me” signals, and the characteristics of infiltrating immune cells in the tumor microenvironment. However, our findings indicate that CD24 does not play a universal role as a macrophage immune checkpoint and that surface expression of CD24 alone does not guarantee its function in this regard.

Our studies also indicate that direct targeting of CD24 carries potential risks for toxicity. In our immunocompetent mouse models, we found that CD24 is expressed on the surface of red blood cells. As a consequence, anti-CD24 antibodies resulted in acute, dose-dependent anemia that was lethal in some mice. CD24 expression patterns are different in the human system, where it is predominantly expressed on neutrophils and B cells in the blood. Although this pattern may pose a different risk for acute toxicity, we found that antibodies targeting human CD24 could deplete neutrophils in ex vivo cultures of unfractionated leukocytes and via macrophage phagocytosis assays. These findings suggest that hematologic toxicity should be cautiously monitored in studies evaluating anti-CD24 agents—particularly neutropenia—since this side effect can be life-threatening if severe.

Regardless of whether CD24 functions as a macrophage immune checkpoint, we found that anti-CD24 antibodies can robustly stimulate macrophage phagocytosis and can be further enhanced by CD47 blockade. These findings are notable since not all antigens can be targeted to elicit immune effector functions, and not all antibodies stimulate phagocytosis as a response. Thus, our findings indicate that strategies to leverage CD24 as a target of opsonization may be valuable if, in light of the risks of toxicity, they offer an acceptable therapeutic index. Such strategies could include intratumoral therapeutic administration or other modalities to maximize antibody concentrations within the tumor microenvironment. Nonetheless, even our bispecific antibody engineering efforts indicate that CD47 may be a better target than CD24 in a bispecific format. Bispecific antibodies that target CD47 with low affinity may achieve maximal macrophage activation while enhancing specificity to the tumor microenvironment. These antibodies warrant further investigation as therapeutics for patients with cancer. Overall, our study provides insight into the relative contributions of CD47 and CD24 in regulating anti-tumor immunity by macrophages, informs the best therapeutic strategies for targeting these molecules, and identifies strategies to overcome liabilities. We expect these findings will further advance the field of macrophage immune checkpoints and will provide insight into the design of better macrophage-directed therapies for cancer.

## METHODS

### Cell lines and culture

KPCA.C (Weinberg lab) was cultured in DMEM (Thermo Fisher) with 10% heat inactivated ultra-low IgG FBS (Thermo Fisher), 1x penicillin/streptomycin/glutamine (Gibco), 1x insulin-transferrin-selenium (Gibco), and 0.2% human epidermal growth factor (25). 238N1 (J. Weissman lab), MC38 (Koch Institute), TUBO-EGFR (I. Weissman lab, Stanford University), 3LL ΔNRAS (J. Downward lab, Francis Crick Institute), and EL4 (I. Weissman Lab, Stanford University) were cultured in DMEM (Thermo Fisher) with 10% heat inactivated ultra low-IgG FBS (Thermo Fisher), 1x penicillin/streptomycin/glutamine (Gibco). NCI-H358, PC9, NCI-H3122 (A. Hata lab, MGH Center for Molecular Therapeutics), KP1 (I. Weissman lab, Stanford University), DLD-1 (I. Weissman lab, Stanford University), control and knockout variants of DLD-1 (Synthego), control and knockout variants of A375 (Synthego), and unmodified A375 (Koch Institute) were cultured in RPMI (Thermo Fisher) with 10% heat inactivated ultra low-IgG FBS (Thermo Fisher), 1x penicillin/streptomycin/glutamine (Gibco). Control and knockout variants of NIH-OVCAR3 (Synthego) were cultured in RPMI (Thermo Fisher) with 20% heat inactivated ultra low-IgG FBS (Thermo Fisher), 1x penicillin/streptomycin/glutamine (Gibco).

### Bone-marrow derived mouse macrophage generation

Primary mouse macrophages were derived from bone marrow as previously described(18,26). Briefly, forelimb and hind leg bones were collected from C57BL/6J or B6129SF1/J mice (Jackson Labs), and bone marrow was extracted using a mortar and pestle. Unfractionated bone marrow was washed with PBS (Thermo Fisher). Red blood cells were lysed using ACK Lysis Buffer (Thermo Fisher) and washed with PBS. Cells were cultured on Petri dishes (Corning) in IMDM (Gibco) containing 10% heat inactivated ultra-low IgG FBS, 1x penicillin/streptomycin/glutamine (Gibco) with 20 ng/ml murine M-CSF (Peprotech) for 7 days. Macrophages were collected for further experimental analysis by incubating with TrypLE Express Enzyme (1X) no phenol red (Thermo Fisher) followed by cell lifting and replating as needed. Macrophages were generally used for experiments between days 7-21 of culture.

### Human macrophage derivation

Primary human macrophages were differentiated ex vivo as previously described(17,27). Briefly, leukocyte reduction system chambers containing heparinized blood were obtained from discarded specimens of anonymous blood donors (Crimson Core Biobank, Brigham and Women’s Hospital). Monocytes were isolated using StraightFrom Whole Blood CD14+ microbeads (Miltenyi) with whole blood purification performed using an autoMACS Pro Separator or autoMACS NEO Separator (Miltenyi). Purified CD14+ monocytes were cultured in IMDM (Gibco) containing 10% heat inactivated ultra-low IgG FBS with 1x penicillin/streptomycin/glutamine (Gibco) and 20 ng/mL human M-CSF (Peprotech) for 7 days. Macrophages were collected for further experimental analysis by incubating with TrypLE Express Enzyme (1X) no phenol red (Thermo Fisher) followed by cell lifting and replating as needed. Macrophages were generally used for experiments between days 7-21 of culture.

### In vitro phagocytosis assays

Cancer cells were collected, washed with PBS and removed using TrypLE, then fluorescently labeled with CellTrace CFSE Cell Proliferation Kit for flow cytometry (Thermo Fisher). Macrophages were collected as described above. Cancer cells and macrophages were combined at a ratio of 4:1 in each well of a ultra-low cluster U-bottom 96-well plate (Corning 7007) in IMDM (Gibco 12440061) with the indicated therapeutic antibodies.

The co-culture was then incubated for 2 hours at 37°C in a humidified incubator with 5% carbon dioxide. At the end of the incubation, cells were washed with cold autoMACS Running Buffer (Miltenyi), and for human experiments stained with a fluorophore-conjugated anti-human CD45 antibody (BioLegend) to identify the macrophages. For murine experiments, cells were stained with a fluorophore-conjugated anti-mouse CD45 or F4/80 antibody (BioLegend) to identify macrophages. Cells were stained with DAPI to exclude dead cells. Phagocytosis was quantified as the percentage of macrophages that had engulfed CFSE-positive cancer cells. All therapeutic antibodies were used at a final concentration of 10 ug/ml.

### Long-term co-culture assays

Cancer cells and macrophages were collected as described above. Macrophages, antibodies, then cancer cells were plated sequentially in 20 ul each of IMDM, no phenol red (Gibco 21056023) with 10% heat inactivated ultra-low IgG FBS (Gibco 16-250-078), 1% penicillin-streptomycin-glutamine (100X) (Gibco 10-378-016), and 20 ng/ml recombinant human M-CSF (PeproTech 300-25) or recombinant murine M-CSF (PeproTech 315-02) into a Corning 384-well microplate (Corning CLS3764). Monoclonal antibodies were used at a final concentration of 10 ug/ml. Plates were allowed to sit at room temperature for 15 minutes before being transferred to an Incucyte S3 Live-Cell Analysis System (Sartorius) in a humidified incubator with 5% carbon dioxide maintained at 37° C. Whole-well images of phase contrast, green fluorescence, and/or red fluorescence were acquired every four to eight hours for up to 7 days using a 4X objective. The growth of cancer cells was analyzed by automated image analysis using the Incucyte Base Analysis Software (Sartorius) to quantify the fluorescent area per well as previously described(17). Statistical comparisons were generally performed at t = 6.5 days as a reference endpoint unless otherwise indicated.

### Lentiviral production and transduction

GFP-luciferase+ lines were transduced by plating 75,000 cells in their respective medium in a 12-well plate (Corning) with 5 uL CMV-GFP-T2A-Luciferase lentivirus (Systems Biosciences) and 8 ug/mL polybrene (Milipore Sigma). Medium was changed after 3 days to allow cells to recover. GFP+ populations were sorted at least two independent times on a BDFACS Aria II machine for GFP+ purity. For generation of CRISPRi-mCherry lines, plasmid pJB109_pHR-UCOE-EF1a-Zim3-NLS-dCas9-HA-2xNLS-P2A-mCherry (28) was co-transfected with delta8.2 and VSV-G into HEK-293T to make CRISPRi-mCherry lentivirus. Stable cell lines expressing CRISPRi-mCherry were generated by infecting cells with the CRISPRi-mCherry lentivirus with 8 ug/ml polybrene. mCherry+ cells were purified by FACS and then confirmed by post-sort analysis of mCherry expression. Lentivirus for mouse or human CD24 overexpression was produced by VectorBuilder using the pLV expression vector with an EF1A promoter driving mCd24a (NM_009846.2) or human CD24 (NM_001291738.1) expression and a puromycin resistance gene. CRISPRi sgRNA, control, and CD24 overexpression lines were created by resuspending 500,000 cells in 1 mL virus with 8 ug/ml polybrene. Medium was changed after 24 hours and cells were allowed to recover before subjecting to puromycin selection. Cells were maintained and passaged in 3 ug/ml puromycin (Selleckchem) after establishment of the lines, and cells were generally used for experiments after at least 5 days of selection. Lentivirus for StayGold (29) expression was produced by VectorBuilder using the pLV expression vector with an EF1A promoter driving hStayGold (Genbank LC593679.1) and contained a puromycin resistance gene. Transduced cells were selected for StayGold expression using puromycin.

### RNP transfection

Knockout cell lines were made using Synthego Gene Knockout Kit v2 and according to the Synthego Immortalized Cell Lipofection Protocol in 24-well plates. In brief, sgRNA and recombinant Cas9 were diluted in OptiMem, and Cas9fectamine was diluted in Optimem in a separate tube. The tubes were mixed together and incubated at room temperature for 5-10 minutes to allow RNP complexes to form. RNP complexes were then added to cells in culture. Medium was changed after 2 days. Cells were passaged and upscaled for further selection by cell sorting. Fluorophore-conjugated antibodies were used to stain and sort for polyclonal populations of cells that exhibited loss of cell-surface CD24 or CD47 protein expression. Cells were sorted at least twice to achieve homogeneous loss of cell-surface expression.

### CRISPR screen

Genome-wide CRISPR screens were performed using the murine lung adenocarcinoma cell line 238N1. Cancer cells were transduced with the mouse Gouda genome-wide sgRNA library (30) at low multiplicity of infection (MOI∼0.3) and selected with puromycin to ensure each cancer cell carries a single and unique sgRNA. Puromycin-selected cancer cells were subjected to 3 distinct conditions: (i) cancer cells alone, (ii) cancer cells and macrophages, (iii) cancer cells, macrophages, and an anti-CD24 antibody. Cancer cells treated with macrophages were co-cultured with primary mouse bone marrow-derived macrophages (BMDMs) at 1:4 ratio for 4 days. Cancer cells were treated with anti-CD24 antibody clone M1/69 at a concentration of 10 ug/mL. Cells were grown at minimum library coverage of 500x for the screen. Genomic DNA from each sample was extracted, the cassette encoding the sgRNA was amplified by PCR, and relative sgRNA abundance was determined by next generation sequencing as previously described(30).

### CRISRPR screen analysis

Sequencing FASTQ files from CRISPR screens were aligned, processed, and counted using custom Python-based scripts (based on https://github.com/mhorlbeck/ScreenProcessing), as previously described(31,32). sgRNA-level phenotypes were normalized as log_2_(enrichment) between sample conditions. Gene-level phenotypes were scored based on the average sgRNA-level phenotype of the 2 sgRNAs targeting a gene. Mann-Whitney test p-values were calculated by comparing all sgRNAs targeting a given gene to the full set of negative control sgRNAs. Discovery scores were calculated by the absolute value of gene-level log_2_(enrichment) over the standard deviation of all evaluated phenotypes multiplied by the −log_10_(p-value). Screen hits were defined as genes with a discovery score greater than 5 (such that FDR is < 0.05). All analyses were performed in Python.

### CRISPR screen Gene Set Enrichment Analysis

Gene set enrichment analysis (GSEA) was performed using FGSEA from the R Bioconductor Suite (v.3.18). Gene Ontology (GO) Biological Process and KEGG pathways were used for analysis. GPI-AP related genes were identified based on previous studies(33).

### Therapeutic antibodies

Antibodies used for experiments included: InVivoMAb anti-mouse/human/rat CD47 (IAP) clone MIAP410 (BioXCell BE0283), InVivoMAb anti-mouse CD24 clone M1/69 (BioXCell BE0360), InVivoMAb anti-human CD47 clone B6.H12 (BioXCell BE0019-1), anti-human CD24 clone ML5 (Biolegend 311102), anti-human CD24 clone SN3 (GeneTex GTX74945), cetuximab (Selleckchem A2000). For FcR blocking experiments, we used anti-mouse Truestain clone 93 (BioLegend 101320) and anti-mouse CD16/CD32 clone 2.4G2 (BioXCell BE0307).

### Syngeneic tumor models

C57BL/6J (Jackson Laboratory #000664) sex and age matched mice were engrafted with appropriate cancer cell lines. Mice were used for engraftment when they were generally 6-12 weeks of age. We used 1 million cells in 100 uL of sterile PBS per subcutaneous engraftment. 238N1 cancer cells were engrafted into B6129SF1/J mice (Jackson Laboratory #101043) at 1 million cells in 100 uL sterile PBS subcutaneously. Tumor dimensions were measured by caliper twice per week and used to calculate tumor volumes according to an ellipsoid formula: length x width x width x π/6. For treatment experiments, therapeutic antibodies were administered by intraperitoneal injection in sterile PBS three times per week or as otherwise indicated. For an intraperitoneal model, 1 million KPCA.C cells were engrafted into the peritoneal cavity in 100 uL sterile PBS. Mice were euthanized according to humane experimental endpoints including tumor size and body conditioning scores.

### Xenograft tumor models

NOD.Cg-Prkdc^scid^ Il2rg^tm1Wjl^/SzJ (NSG) (Jackson Laboratory #005557) sex and age matched male or female mice were engraftment subcutaneously with cancer cells in a 1:1 ratio of PBS and Matrigel Matrix hESC-Qualified Matrix (Corning 354277). Mice were used for engraftment when they were approximately 6-12 weeks of age. Mice were engrafted with 1 million cancer cells in 100 uL sterile PBS subcutaneously. Tumor volumes were measured and calculated as described above. Mice were euthanized according to human experimental endpoints including tumor size and body conditioning scores.

### Mouse blood analysis

Venous blood samples from mice were collected via retro-orbital sampling into Microvette 100 EDTA tubes (Sarstedt). Blood was analyzed for hematologic parameters by IDEXX CBC (MIT DCM Diagnostics Lab). Receptor occupancy was examined by flow cytometry using an anti-mouse IgG secondary antibody conjugated to AlexaFluor647 (BioLegend). Red blood cells were prepared for in vitro phagocytosis assays by CFSE-labeling, then the cells were co-cultured with primary mouse macrophages as described above. For quantification of internalization of red blood cells, red blood cells were labeled with pHrodo (Thermo Fisher) according to manufacturer’s instructions. Blood cells were then incubated with primary mouse macrophages and imaged using a Incucyte S3 Live Imaging System (Sartorius). pHrodo-positive macrophages were quantified over time and evaluated as a measure of phagocytosis using Incucyte Base Analysis Software (Sartorius).

### Human blood analysis

Unfractionated blood from LRS chambers containing anticoagulant, or EDTA-anticoagulated peripheral blood (Miltenyi Biotec) was used for analysis. Blood specimens were depleted of red blood cells using EasySep RBC Depletion Reagent and “The Big Easy” EasySep Magnet according to the manufacturer’s protocol (STEMCELL Technologies). For viability assays, cells were then counted and plated in ultra-low retention 96-well U-bottom plates (Corning #7007) at 200k cells per well in IMDM medium containing 10% heat-inactivated ultra-low IgG FBS and 1x pen/strep/glutamine and 10 ug/ml of the indicated antibodies. Cells were incubated at 37°C for 24 hours and stained for the indicated markers and assessed for viability using a BD LSRFortessa Cell Analyzer (BD Biosciences). For neutrophil phagocytosis assays, primary neutrophils were isolated from RBC depleted blood using MACSxpress Whole Blood Neutrophil Isolation Kit, human (Miltenyi Biotec) and purified according to the manufacturer’s protocol. Neutrophils were labeled using CellTrace CFSE Cell Proliferation Kit (Thermo Fisher). 200,000 neutrophils were co-cultured with 50,000 primary human macrophages with 10 ug/ml of the indicated antibodies as appropriate for 2 hours at 37°C in 100 ul serum-free IMDM. Cells were then stained with APC anti-human CD45 as previously described and analyzed on a BD LSRFortessa Cell Analyzer. Macrophages were distinguished from neutrophils based on forward and side scatter as well as high expression of CD45. Phagocytosis was quantified as the percentage of macrophages that had engulfed CFSE+ neutrophils.

### Cytokine Analysis

Cell culture supernatants from unfractionated leukocytes were collected after 24 hours of culture by centrifugation followed by freezing at −80 °C. Supernatants were subjected to analysis using a Human Cytokine 71-Plex Discovery Assay (Eve Technologies). Data were analyzed by creating a matrix of observed cytokine concentrations, setting any out-of-range measurements to the maximum observed level of that cytokine (if above the calibration range) or 0 (if below). Cytokines were dropped from the analysis if they were within range in at most one condition. Cytokine levels were compared to the control condition using one-way ANOVA followed by Dunnett’s method for multiple comparisons. To generate a heatmap of cytokine levels relative to the control mean, cytokines were clustered by uncentered correlation, with clusters merged by pairwise average-linkage.

### Flow Cytometry

Cells were stained in autoMACS Running Buffer (Miltenyi Biotec 130-091-221) at the recommended concentrations and were analyzed using a BD LSRFortessa Cell Analyzer. When necessary, anti-mouse or anti-human TrueStain (Biolegend) was used to block Fc receptors. In general, cells were stained on ice for 30 minutes with primary antibodies. Cells were then washed twice and resuspended in 100 ng/mL DAPI (Millipore-Sigma). Antibodies used include APC anti-mouse CD24 (BioLegend 101814), APC anti-mouse CD47 (BioLegend 127513), APC anti-mouse CD45 (BioLegend 103112), AlexFluor647 anti-mouse F4/80 (Biolegend), APC anti-human CD45 (Biolegend), APC anti-human CD24 (Biolegend), and APC anti-human CD47 (Biolegend). As appropriate, compensation was performed using Ultracomp beads(BD). Analysis was generally performed by excluding debris and doublets based on forward and side scatter and excluding dead cells based on DAPI staining. Analysis of geometric mean fluorescence intensity and population percentages were performed using FlowJo version versions 9-10 (TreeStar).

### Single-cell RNA Sequencing Sample Preparation

Tumors were obtained from euthanized mice and put into 6-well plates (Corning) with PBS. Tumors were weighed and placed into gentleMACS C tubes (Miltenyi Biotech) containing RPMI (Gibco) with 1% collagenase (Sigma Aldrich) and 1% hyalunorase (Sigma Aldrich) on ice. Tumors were then dissociated in an OctoDissociator (Miltenyi Biotech) for 1 minute. The tubes were then incubated at 37°C for 30 minutes on a shaker. Homogenized cell suspensions were filtered through 70 um (Falcon), then 40 um (Falcon) cell strainers and centrifuged. The supernatant was removed and RBCs lysed using ACK lysis buffer (Thermo Fisher) for 1 minute. Cells were centrifuged again and stained with anti-mouse CD45.2 APC (Biolegend) or anti-human CD45 APC (Biolegend) for 20 minutes on ice. Cells were centrifuged and resuspended in DAPI (Sigma Aldrich) and sorted for DAPI negative APC positive cells. Sorted cells were centrifuged and aspirated, then resuspended in cell staining buffer (Biolegend) with anti-mouse or human TrueStain and incubated on ice for 10 minutes. To each cell population, 1 µg hashtagging antibody (Biolegend) was added and cells were incubated on ice for 30 min. Cells were washed with Cell Staining Buffer twice and counted. Cells were brought to 1000 cells/µL and pooled appropriately for sequencing.

### 10x Library Preparation and Sequencing

Cells were processed using the 10X Genomics Chromium Controller with the Single Cell 3ʹ v3.1 Reagent Kit, according to manufacturer’s directions. Briefly, a target number of cells per library was roughly achieved by loading 1.65 times the target number of cells in suspension, along with barcoded beads and partitioning oil, into the Chromium Controller, in order to create GEMs (Gel Beads in Emulsion). The Chromium Controller combines individual cells, first strand master mix, and gel beads containing barcoded oligonucleotides into single-cell droplets for first strand cDNA synthesis, so that each cell is marked with its own unique barcode during reverse transcription. The 3’ beads contain a poly(dT) oligo that enables the production of barcoded, full-length cDNA from poly-adenylated mRNA. After first strand synthesis is complete, the emulsion is dissolved and the cDNA is pooled for bulk processing as a single sample. The sample is fragmented, end-repaired, A-tailed and ligated with universal adapters. A second sample barcode is then added during the PCR step, allowing for unique library identification. The end result is a single library representing one cell suspension, containing data for each individual cell. qPCR was performed on all libraries using KAPA qPCR library quant kit as per manufacturer’s protocol. The samples are loaded on the NovaSeq 6000 based on qPCR concentrations. The demultiplexing and fastq generation is performed using illumina’s BCL to FASTQ file converter bcl2fastq v2.20.0.422.

### Single-cell RNA Sequencing Analysis

The same processing and analysis pipeline was used across all samples and all batches. Raw read processing was performed using the 10X Genomics workflow. We used the CellRanger Single-Cell Software Suite (v7.0.0) to perform barcode assignment and unique molecular identifier (UMI) quantification. The reads were aligned to the 10X reference genome mm10 (ref-2020-A). Seurat package (v5.1.0) was used for demultiplexing cells based on TotalSeq barcodes. Any cells which were not associated with TotalSeq barcode (negative) or assigned to more than 1 TotalSeq barcode (doublet) were discarded. Data for all singlet cells were normalized and scaled using default parameters and then integrated using "HarmonyIntegration" method implemented in seurat. We use the AUCell bioconductor package to identify immune specific marker sets (defined in Panglaodb - https://panglaodb.se/) that are highly expressed in each cell for cell type determination. FindMarkers function implemented in seurat was used for identifying differentially expressed genes. We also perform a functional Gene Set Enrichment Analysis (GSEA) for all genes using hallmark genesets defined in msigDB(https://www.gsea-msigdb.org/gsea/msigdb/) and the fgsea R package.

### Plasmid Preparation for scFv Library

Antibody and binding domain sequences were curated from publicly available databases and literature sources and included the following: EGFR (cetuximab, IMGT 7906), EpCAM (US Patent No.: US 9,777,073 B2), HER2 (trastuzumab, IMGT 1n8z), PD-L1 (atezolizumab, KEGG D10773), TROP2 (sacituzumab, KEGG D10984), FOLR1 (mirvetuximab, KEGG D10953), Nectin-4 (enfortumab, KEGG D11524), Netrin-1 (NP137(34)), CD71 (delpacibart, IMGT 1374), CV1(18), WTa2d1(18), SIRPa (KWAR23(35)), CD24-1(36), CD24-2 (US Patent Application Pub. No.: 20210213055A1), CD24-3 (US Patent No.: US 8,614,301 B2), PD-1 (nivolumab, KEGG D10316). The sequences were reverse-translated, codon-optimized for *Homo sapiens* and then engineered with flanking 5’ and 3’ cloning sites. Gene fragments containing the sequences were obtained from Twist Biosciences and were then cloned into a modified pFuse vector (Invivogen) containing a human IgG1 knob or hole construct. Cloning was performed using New England Biolabs restriction enzymes for DNA digestion followed by T4 DNA ligase fusion and bacterial transformation with chemically competent *E.coli* TOP10 cells. Transformed cells were selected on LB agar plates containing 40 μg/ml zeocin. Single colonies were then used to inoculate 5 mL of LB medium containing the same concentration of antibiotic. The culture was used for plasmid purification using the ZymoPure Plasmid Purification kit following manufacturer instructions. The concentration of the plasmid product was assessed using a ThermoFisher Nanodrop by taking the absorbance at 260 nm and the correct insertion was verified by whole plasmid DNA sequencing (Quintara Biosciences).

### Expression and Purification of Recombinant Antibodies

Purified pairs of IgG1-Knob and IgG1-Hole plasmids were co-transfected to a final concentration of 1 ug/ml of each into 900 μl of Expi293F cells at 3 × 10 cells/ml (Thermo Fisher Scientific, A14528) in 96 deep-well plate (USA Scientific, 1896–2110) with ExpiFectamine 293 Transfection Kit (Thermo Fisher Scientific, A14524) following the manufacturer’s recommendation and incubated at 37°C, 8% CO_2_ with shaking at 900 rpm for 7 days. The antibody supernatants were diluted 2-fold with PBS and used for further analysis.

### Research Ethics Statement

The animal experiments performed in this study were approved by the Massachusetts Institute of Technology Committee on Animal Care. This research has been conducted following ethical standards related to research involving animals including the use of humane experimental endpoints.

## ACKNOWLEDGEMENTS

We thank members of the Weiskopf lab as well as members of the Tobiloba Oni and Robert Weinberg labs for experimental assistance, reagents, and insightful discussions. We thank Donna Hicks, Elinor Eaton, Joana Liu Donaher, Nicholas Polizzi, and Beverly Dobson for support. We thank Irving Weissman and Julian Downward for providing cell lines and reagents. We thank Patrick Autissier and the Whitehead Institute Flow Cytometry Core Facility; Stephen Mraz, Thomas Vokert, and the Genome Technology Core Facility; and the Bioinformatics and Research Computing Core Facility. Experimental diagrams were created using Biorender. This work was supported by the Valhalla Foundation (to KW); NIH grant T32 CA09172 (to KW); A Breath of Hope Lung Foundation Peg Fisher-Jullie Fight for Life Award (to KW); ASCO Conquer Cancer Foundation Young Investigator Award (to KW); AACR-AstraZeneca Career Development Award for Physician-Scientists in Honor of José Baselga (to KW); Fast Grants, Emergent Ventures, George Mason University (to KW); Society for Immunotherapy of Cancer, Holbrook Kohrt Cancer Immunotherapy Translational Memorial Fellowship (to KW); American Lung Association Catalyst Award (to KW); an anonymous grant (to KW); Richard Reisman in honor of Jane Reisman and Lilian Reisman (to KW); Department of Defense grant W81XWH2210141 (to KW); NIH grant R01 CA279259 (to KW); Research Foundation for the Treatment of Ovarian Cancer (to KW); National Cancer Institute Cancer Center Support Grant P30-CA014051 (Koch Institute High-Throughput Sciences Facility). D.Y. is supported by a Damon Runyon Dale Frey Award, an NCI Transition Career Development Award 1K22CA289207 and an NIH Director’s New Innovator Award 1DP2OD037078. The content of this manuscript is solely the responsibility of the authors and does not necessarily represent the official views of the NIH or other funding agencies.

## AUTHOR CONTRIBUTIONS

JA, AM, KV, JV, JR, JB, RM, KW designed and performed co-culture experiments in vitro. KV, VC, RH, JC, JW, DY, KW designed, performed, and/or analyzed CRISPR screening experiments. JA, AM, KV, FR, KW designed and performed mouse experiments. JA, AM, SG, JV, JL, KW designed, performed, and/or analyzed scRNA-seq experiments. JA, AM, GWB and KW performed and/or analyzed cytokine experiments. JR and KW designed experiments and analyzed data pertaining to bispecific antibodies. RW, DY, JW, KW performed experimental oversight and funding acquisition. JA, AM, KW wrote the first draft of the manuscript, which was subsequently reviewed and/or edited by all co-authors.

---

**Supplementary Figure S1:**
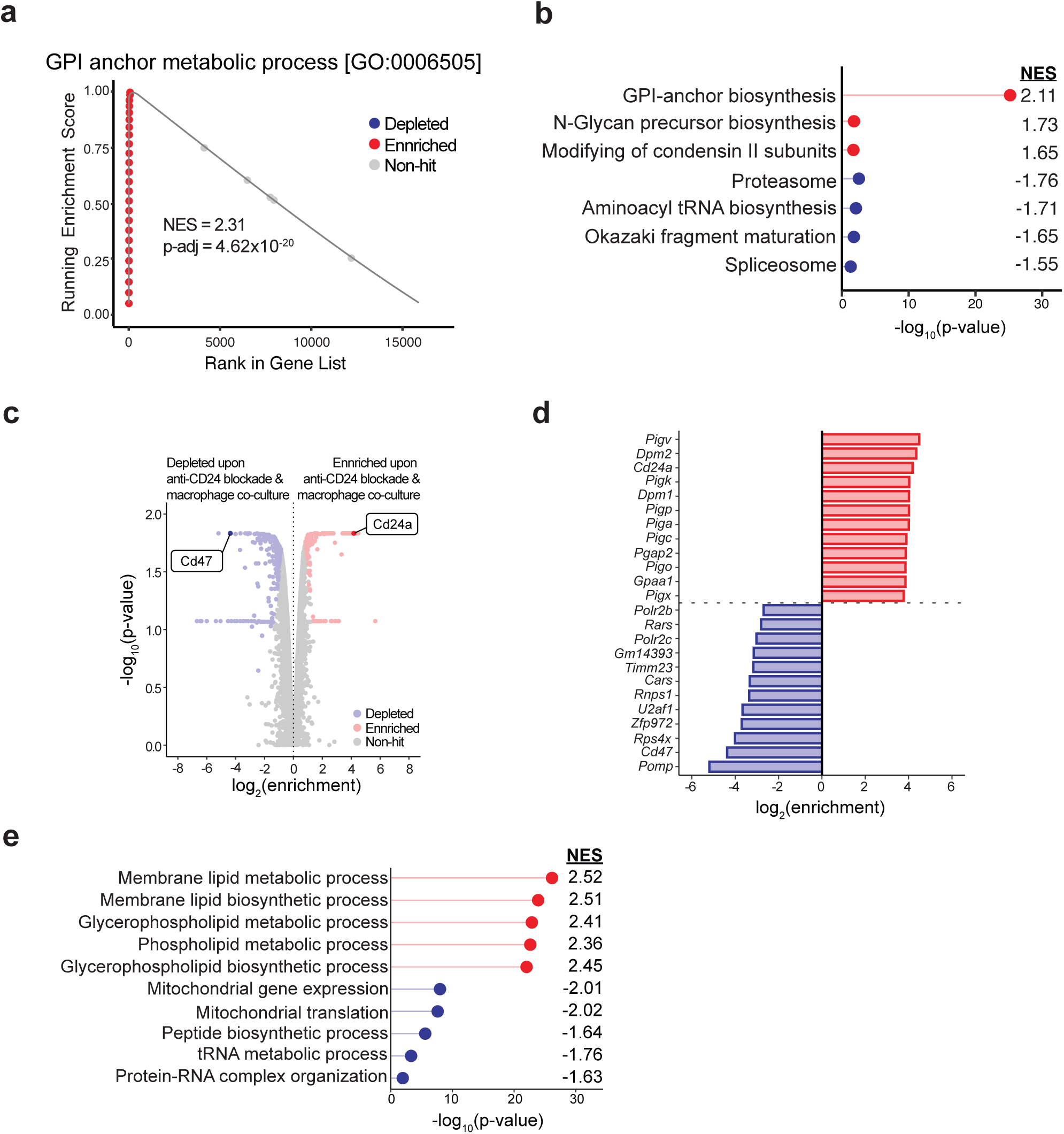
CRISPR screen differentiates immune checkpoint function of CD47 and CD24 in vitro. **A,** Enrichment plot of GPI anchor metabolic process (GO Biological Process) on genes differentially enriched in treatment with macrophages and anti-CD24 against treatment with macrophages alone. Enriched genes are shown in red, and depleted genes are shown in blue. **B,** Differentially enriched KEGG pathways comparing treatment with macrophages and anti-CD24 against treatment with macrophages alone (p-adjusted <0.05). **C,** Volcano plot of genome-wide screen comparing treatment with macrophages and anti-CD24 against cancer cells only. Genes with a discovery score > 5 (FDR < 0.05) are highlighted, with enriched genes shown in red and depleted genes shown in blue. **D,** Top differentially enriched and depleted genes with respective log_2_(enrichment) from differential enrichment analysis depicted in **C. E,** Differentially enriched GO Biological Process pathways comparing treatment with macrophages and anti-CD24 against cancer cells only. Top 5 enriched and depleted pathways are shown (p-adjusted <0.05).

**Supplementary Figure S2:**
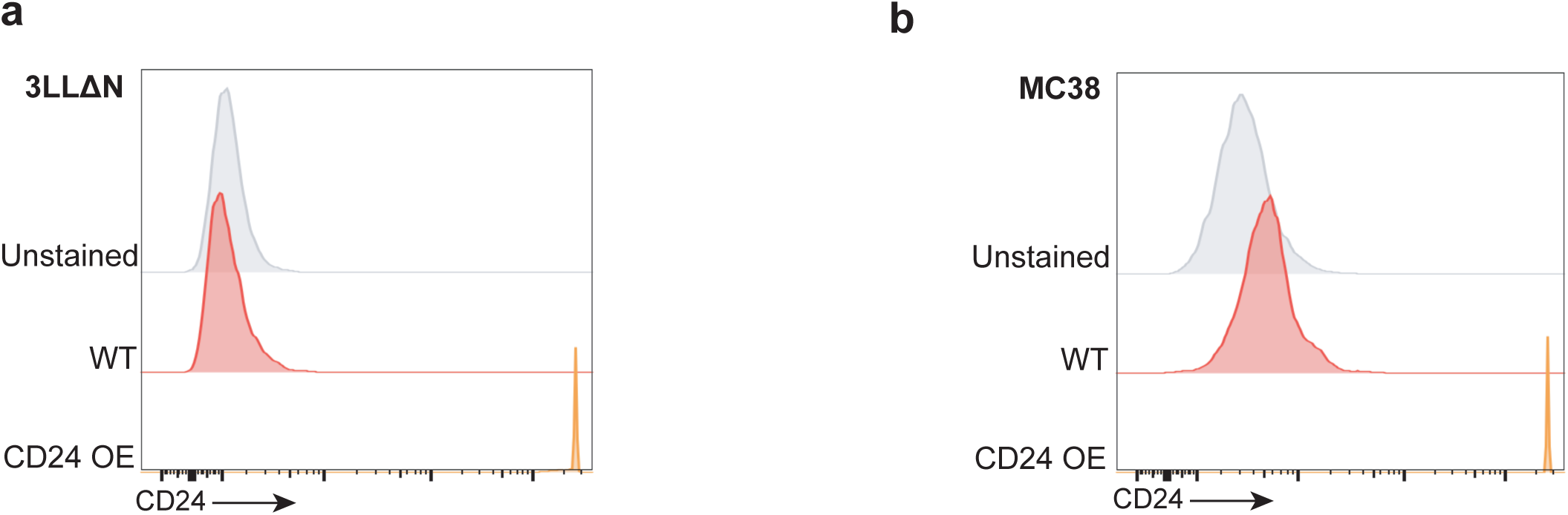
Mouse overexpression cell line validation. CD24 expression as measured by flow cytometry of MC38 (**A**) and 3LL ΔNRAS (**B**) CD24 overexpression cell lines.

**Supplementary Figure S3:**
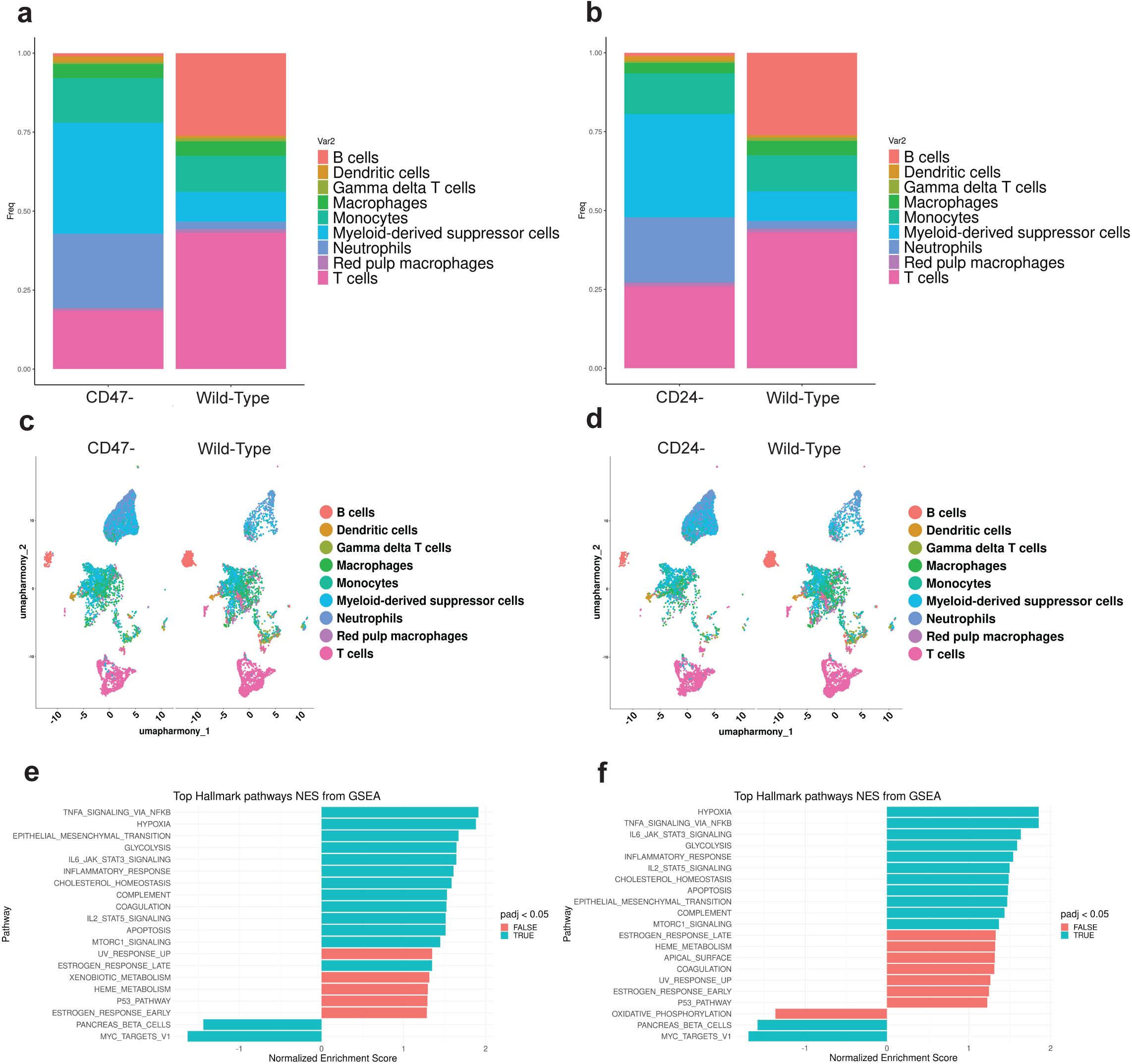
Syngeneic single-cell RNA-sequencing gene enrichment scores. Results of scRNA-seq of sorted CD45+ immune cells from experiments using CD24 or CD47 knockout tumors. (**A,C,E**) Comparison of CD47- tumors (KPCA.C CD47 knockout, 238N1 CD47 knockout) to wild-type tumors (KPCA.C control, 238N1 control). **A,** Relative frequencies of immune cells from CD47- versus wild-type tumors. **C,** UMAP showing identified cell clusters. **E,** Gene set enrichment analysis showing Normalized Enrichment Scores of top Hallmark pathways. (**B,D,F**) Comparison of CD24- tumors (KPCA.C CD24 knockout, 238N1 CD24 knockout) to wild-type tumors (KPCA.C control, 238N1 control). **B,** Relative frequencies of immune cells from CD24- versus wild-type tumors. **D,** UMAP showing identified cell clusters. **F,** Gene set enrichment analysis showing Normalized Enrichment Scores of top Hallmark pathways.

**Supplementary Figure S4:**
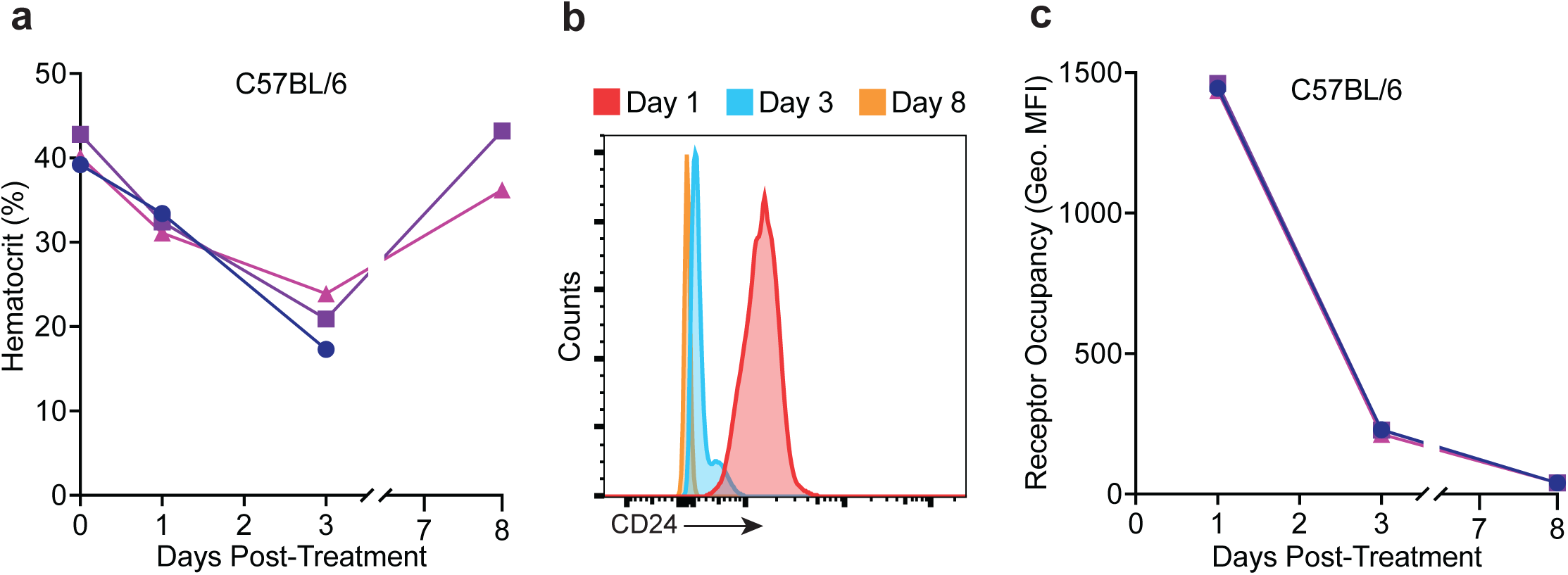
Drop in hematocrit levels upon CD24 targeting in syngeneic model. **A,** C57BL/6J mice were treated with 200 ug anti-mouse CD24 antibody. Hematocrit levels are represented as the mean ± SEM of 3 mice measured at the indicated days post-treatment. **B,** Representative histogram depicting receptor occupancy of anti-mouse CD24 antibody on the surface of mouse red blood cells as detected by staining for antibody binding at the indicated days post treatment. **C,** Quantification of receptor occupancy of anti-CD24 on the surface of red blood cells. Data represents mean ± SEM of n = 3 mice.

**Supplementary Figure S5:**
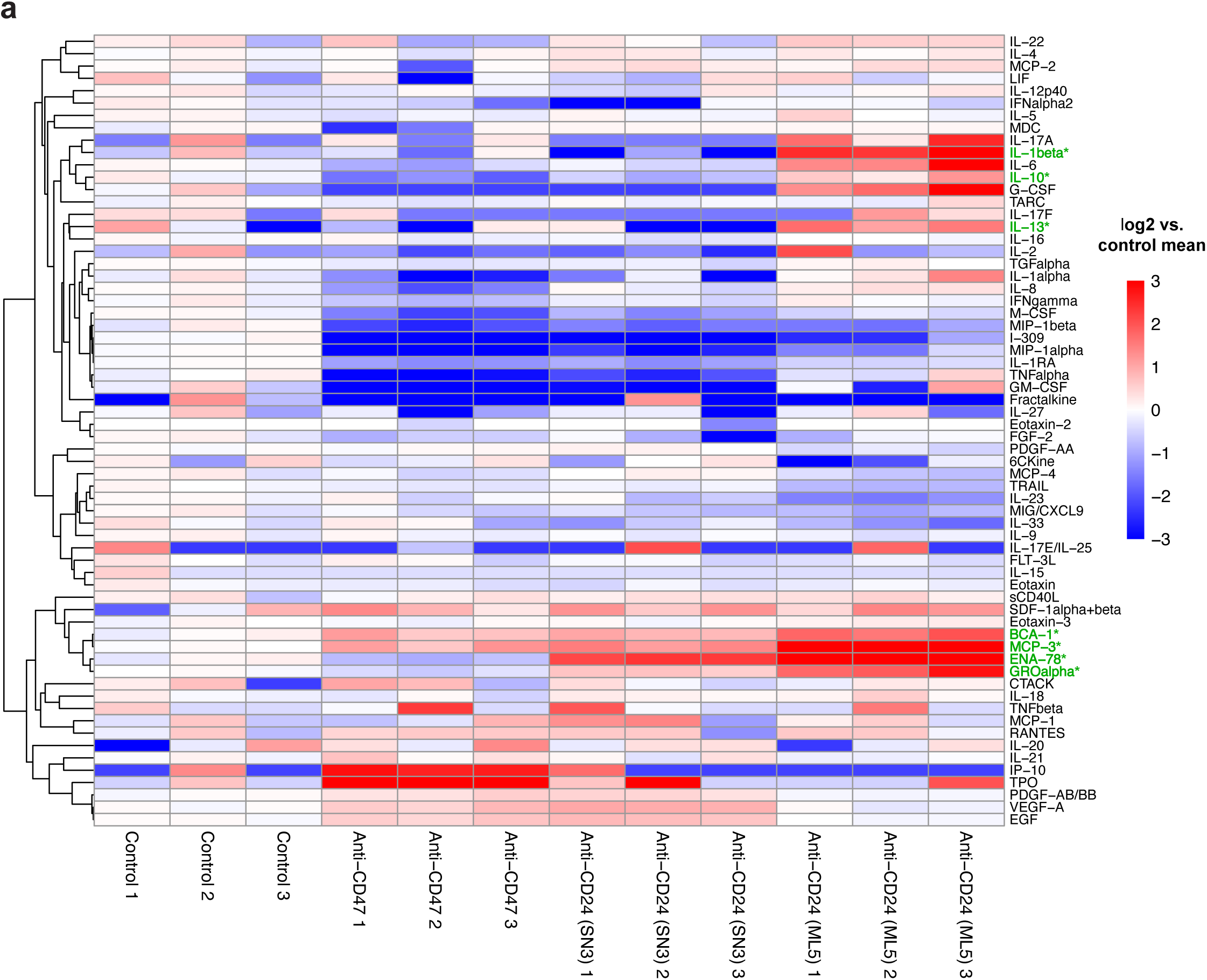
Depletion of neutrophils by anti-CD24 corresponds to secretion of proinflammatory cytokines. **A,** Heatmap of cytokine secretion from unfractionated leukocytes 16 hours after culture with the indicated antibodies utilizing a Human Cytokine 71-Plex Discovery Assay. Cytokines in blue indicate a depletion from the control supernatant while cytokines in red indicate enrichment compared to control supernatant. Cytokines highlighted in green differed significantly between control and anti-CD24 (ML5) treatment.

**Supplementary Figure S6:**
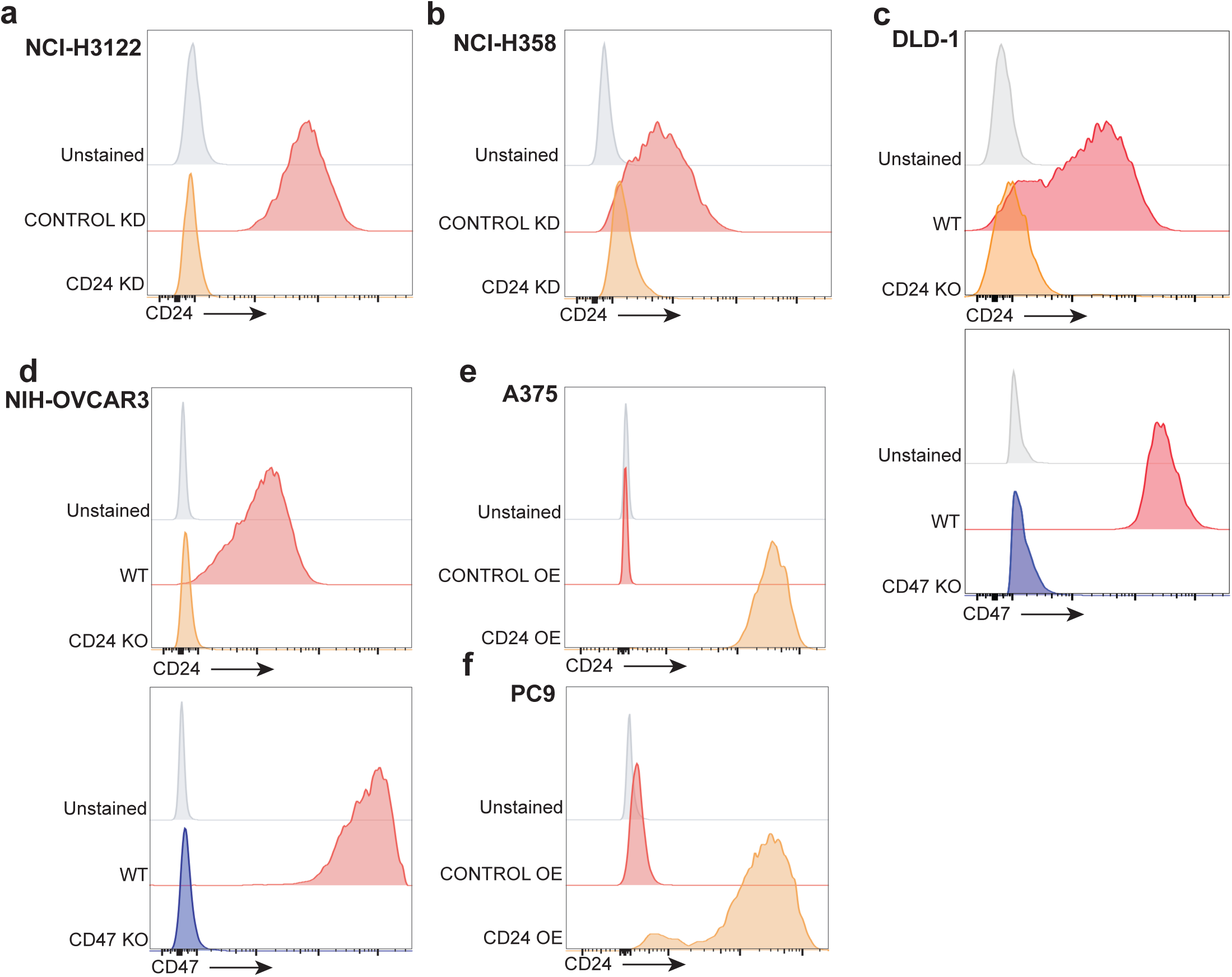
Human knockdown, knockout, and overexpression cell line validation. CD24 expression of CD24 knockdown cell lines (**A**) NCI-H3122, (**B**) NCI-H358,and knockout cell lines (**C**) DLD-1, (**D**) NIH-OVCAR3, and overexpression cell lines (**E**) A375 and (**F**) PC9 as measured by flow cytometry.

